# Uncovering the flexibility of CDR loops in antibodies and TCRs through large-scale molecular dynamics

**DOI:** 10.1101/2025.11.10.687725

**Authors:** Matteo Cagiada, Fabian C. Spoendlin, King Ifashe, Charlotte M. Deane

## Abstract

Antibody structures are composed of framework regions that adopt a conserved fold and complementarity determining regions (CDR) loops which are far more variable. Flexibility of CDR loops has been linked to key properties such as affinity and specificity. However, owing to the scarcity of available data it has not been possible to study the functional implications of their dynamics in detail. To overcome these data limitations, we introduce CALVADOS 3-Fv, a customised set-up of the residue-based CALVADOS 3 model, utilizing restraints and parameterization tailored for immune receptor simulations. CALVADOS 3-Fv reproduces ensemble metrics in all atom simulations and experimental data with high accuracy. Having validated our protocol, we created FlAbDab and FTCRDab, two databases containing simulations of more than 150,000 antibodies and T-cell receptors. The databases are released open source to enable the study of CDR dynamics and as a large data source for training machine learning models.

## Introduction

Antibodies are a vital component of the adaptive immune system and are also used as biother-aupetics and research tools (Lu et al., 2020; Khetan et al., 2022). They consist of four polypeptide chains, two heavy and two light, with antigen interactions mediated by the most N-terminal domain of each chain: the fragment variable (Fv). The Fv comprises of framework regions which adopt conserved secondary structures and six hyper-variable complementarity determining regions (CDRs, 3 on each chain) that drive antigen binding. While framework regions support antibody stability (Julian et al., 2017), the CDR loops are structurally diverse and potentially dynamic (Kroon et al., 2003; Fernández-Quintero et al., 2019a).

It has been suggested that the natural immune system exploits CDR conformational flexibility to balance between promiscuous (polyspecific), low-affinity binding and specific, high-affinity interactions. Structural diversity of an antibody CDRs may enable a single antibody to recognise a broader range of antigens, facilitating polyspecific and cross-reactive recognition, for example binding of mutated antigens from divergent viral strains or distinct envelope proteins (James et al., 2003; Fernández-Quintero et al., 2019; Guthmiller et al., 2020). Flexibility also influences the thermodynamics of antigen recognition by modulating both the enthalpic and entropic cost of binding (Mikolajek et al., 2022). The observed trend of paratope rigidification upon complex formation (Kroon et al., 2003; Fernández-Quintero et al., 2019a) implies that the entropic penalty can be reduced by a pre-configuration of an optimal binding conformation in solution which, in turn, can lead to faster binding kinetics and higher affinities (Li et al., 2015; Schmidt et al., 2013; Haidar et al., 2014).

In case studies of antibody lineages, affinity maturation was shown to introduce mutations that promote CDR rigidification and thereby narrow specificity whilst enhancing binding strength. (Schmidt et al., 2013; Babor and Kortemme, 2009; Li et al., 2015; James et al., 2003; Blackler et al., 2022; Fernández-Quintero et al., 2019b). Thus, CDR flexibility may support a two-stage strategy: the initial promiscuous binding of a broad range of antigens by germline antibodies, which increases the effective size of the immune repertoire, followed by selection for more specific, high-affinity binders. Although conformational flexibility may be important in modulating functional properties of some antibodies, its functional relevance and applicability at the repertoire level remain unclear (Jeliazkov et al., 2018).

Molecular dynamics (MD) simulations are widely used to study protein dynamics at atomic resolution, and have been applied to study antibody flexibility (Wong et al., 2011; Fernández-Quintero et al., 2019b; Guloglu and Deane, 2023). All-atom MD is particularly well-suited for capturing a range of molecular interactions. However, the high computational cost of all-atom MD limits long-timescale sampling and the number of proteins that can be simulated. Even the largest all-atom simulation datasets, such as ATLAS (Vander Meersche et al., 2024) and mdCATH (Mirarchi et al., 2024), contain a number of simulations far lower than the number of unique experimentally determined structures. Coarse-grained (CG) models offer a more scalable alternative, as they reduce the number of particles in the system and thereby increase sampling efficiency. Recent studies have demonstrated that CG models can efficiently simulate datasets of up to 30,000 proteins (Tesei et al., 2024), offering accurate molecular descriptions within a reasonable computational time. Generative structure prediction models are increasingly being explored for modeling conformational ensembles (Lewis et al., 2025; Zhu et al., 2024; Zheng et al., 2024; Lu et al., 2023; Huguet et al., 2024; Jing et al., 2024; Janson et al., 2025; Invernizzi et al., 2025). Although promising, these approaches currently struggle to accurately capture structural diversity largely due to the limited availability of high-quality data of structural ensembles for training. For antibodies in particularly, performance has not been benchmarked extensively and, therefore, applicability remains uncertain (Spoendlin et al., 2025; Riccabona et al., 2024).

High quality experimentally determined structures is available for more than 3,000 unique antibodies (Dunbar et al., 2014; Schneider et al., 2022). If machine learning-based predictions are also considered the number of available antibody structures is far greater. CG molecular dynamics models offer the most viable approach to extensively investigate the dynamics of a dataset of this size. However, simulating antibody Fvs with CG models comes with several difficulties. Fvs contain alternating potentially flexible loops, e.g. the CDRs, and regions with rigid structures, the framework. A suitable model, therefore, must accurately describe both flexible and “folded” regions. A few such models have been proposed, including different versions of the Martini force field (Benayad et al., 2021; Thomasen et al., 2022, 2023) and more recently CALVADOS 3 a residue-based model specifically designed for multi-domain proteins (Cao et al., 2024; von Bülow et al., 2025b).

In this study, we leverage the efficiency of CALVADOS 3, to simulate and characterise the flexibility of antibody and T-cells receptors (TCR) Fvs. The Fv region of these immunoproteins is an ideal candidate for CALVADOS 3 because it consists of two immunoglobulin domains, each of which has alternating CDR loops (disordered regions) and a layer of seven to nine antiparallel β-strands (folded regions). We optimized our simulation protocol to replicate results obtained from all-atom MD simulations and demonstrate that our setup achieves high accuracy in reproducing ensemble metrics. We were able to characterize the flexibility of 3,140 non-redundant, quality-filtered Fv structures from the SAbDab (Dunbar et al., 2014; Schneider et al., 2022) and 280 TCRs from the STCRDab (Leem et al., 2018).

Due to the low computational cost of this approach (two CPU hours per simulation), we incorporated predicted antibody structures into our dataset and, ultimately, simulated over 150,000 immune receptors. We openly release our simulations as new databases in the form of the Flexibility Antibody Database (FlAbDab) and the Flexibility TCR Database (FTCRDab). These large datasets will to provide a resource to investigate the functional role of CDR conformational flexibility and enable the training of machine learning models on a more information rich representation of antibody structure.

## Results

To assess the structural flexibility of CDR loops, we used CG MD simulations to generate and characterise conformational ensembles of Fvs for over 3000 experimentally resolved and 150,000 computationally predicted antibody structures – (Fig. 1), as well as 280 TCR Fvs.

**Figure 1.**
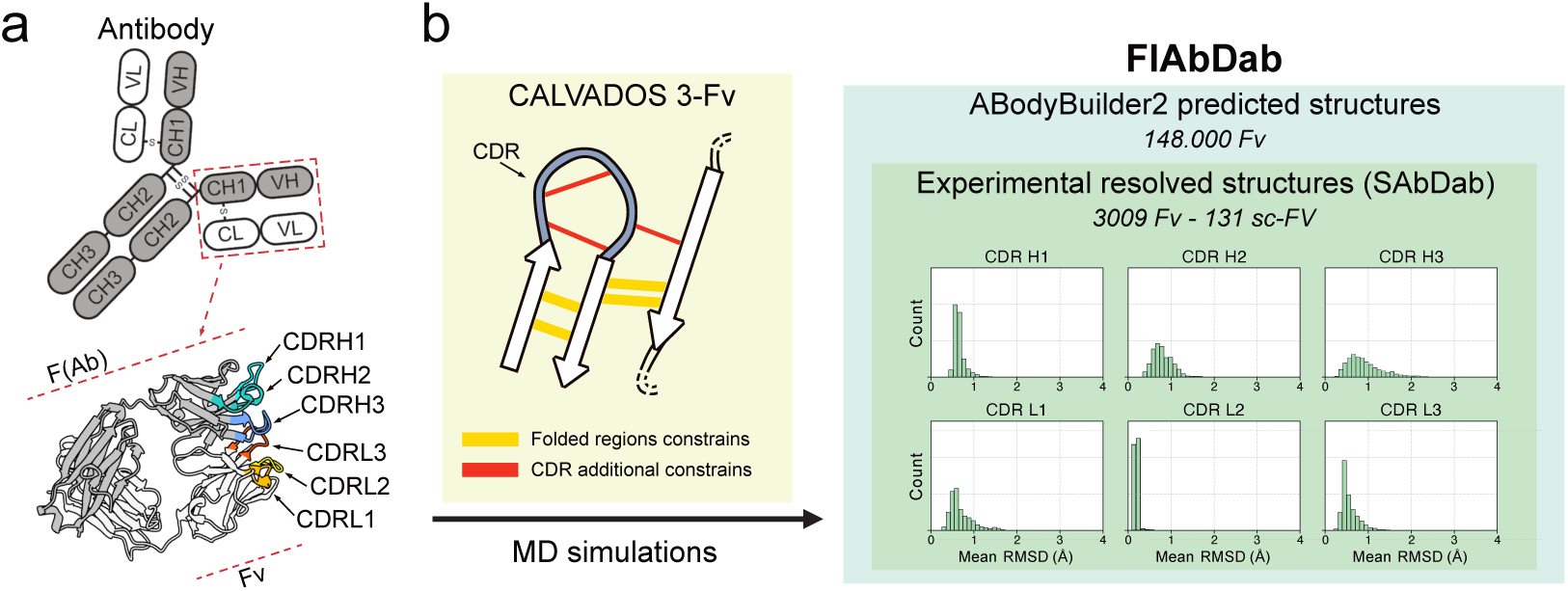
Schematic illustration of our approach to characterise flexibility in antibody Fv regions and generate conformational ensemble using molecular dynamic simulations. (a) We characterised the flexibility of a fragment variable (Fv) region of an antibody, especially looking at complementarity-determining regions (CDRs). (b) We performed molecular dynamics simulations based on CALVADOS 3-Fv, an antibody *ad-hoc* setup for the residue-level model CALVADOS 3. In CALVADOS 3-Fv, additional restraints are introduced to better characterise the flexibility of CDR residues. This enabled more accurate modelling of Fv loop flexibility at a lower computational cost. (c) We used CALVADOS 3-Fv to create our new database, FlAbDab, which can be used to investigate Fv dynamics and train new models. FlAbDab contains over 150,000 simulated Fvs, including those with experimentally solved and computationally predicted structures. We also used CALVADOS 3-Fv simulations to characterise all the unique experimental structures in SAbDab.

### Characterisation of Antibody Fv using CALVADOS 3

We first evaluated the accuracy of the CALVADOS 3 model for simulations of Fv. We simulated Fv regions with CALVADOS 3 by applying harmonic potential to folded regions outside CDR loops and leaving the CDR loops free from additional constraints. The CDR loop boundaries were here defined using IMGT definitions (Lefranc et al., 2005) after the structures were IMGT numbered using ANARCI (Dunbar and Deane, 2016). Hereafter, we refer to this simulation setup as CV3-IMGT. To benchmark the accuracy of the CV3-IMGT setup, we used two different Fvs: the first from a mouse anti-hen egg white lysozyme antibody (F10.6.6, PDB code 2Q76), and the second from an anti-HBV e-antigen monoclonal antibody (anti-HBeAg e6, PDB code 3V6F). For both Fvs, we used as comparison in-house all-atom MD simulations (Fig. S1) and cluster structures from previously published metadynamics (metaD) simulations (Spoendlin et al., 2025). We found that. in this setup, CALVADOS 3 simulations of the Fv tended to overestimate the flexibility of all CDRs in both chains (Fig. S2).

We hypothesised that this excessive flexibility observed was caused by misrepresenting CDR loops as disordered regions. CDR loops often include secondary structure elements (Fig. S3), such as extended β-strands or α-helices (Chothia and Lesk, 1987; Weitzner et al., 2015) and show a significant number of non-covalent interactions, which are typical of folded regions, between their residues as well as with residues in framework regions or in different CDRs. To capture these additional CDR interactions in our CALVADOS 3 simulations, we applied additional constraints to the CDR loops using information extracted from the Fv input crystal structures. We named this setup CALVADOS 3-Fv (CV3-Fv).

We evaluated the accuracy of our new setup by testing it on the two antibody systems used to evaluate the CV3-IMGT setup, as well as two additional TCR systems (PDB 5BS0 and 7F5K). First, we observed an improved agreement in CDR flexibility predictions between our CV3-Fv simulations and all-atom MD simulations, compared to CV3-IMGT simulations (Fig. S2). We also used the harmonic ensemble similarity metric from the ENCORE package (Tiberti et al., 2015) to evaluate the similarity of the CV3-Fv ensembles to the all-atom simulation ensemble. We confirmed that, for all simulated antibody and TCR systems and CDRs, the CV3-Fv setup gave greater agreement with the all-atom simulation than with the CV3-IMGT simulations (Fig. S2e,f, Fig. S12).

We then used ‘frame coverage’, defined as the minimum distance between each all-atom frame and any CV3-Fv frame, to assess the sampling of conformational states (2a,e and S11a,f). In the antibody systems, CV3-Fv simulations captured over 80% of the conformational space observed in the all-atom simulations across all CDR loops within a 1Å threshold. At individual level, frame coverage exceeded 90% for loops L1, H1, L2, H2 and H3; and reaches approximately 60% for H3. We observed similar results for the TCR systems, with high coverage for many of the CDRs, with nearly 100% of frames covered below 1Å by the CV3-Fv protocol. For CDRs A2 and A3 of 5bs0 and B3 of TCR 7f5k, we observed slightly lower values falling to a minimum of 66%. These results, together with the correlations observed in the ensemble similarity metrics described above, suggest that CV3-Fv simulations achieve a close sampling similarity to all-atom simulations.

As conformational coverage alone does not provide insight into the overall flexibility of the ensemble, we looked at commonly used flexibility metrics such as frame-pairwise root-mean-square deviation (RMSD) and root-mean-square fluctuations (RMSF) to collect additional information. We found that the CV3-Fv setup accurately captures the all-atom dynamics of both rigid and flexible loops in antibodies and TCR systems, with RMSD and RMSF values that closely match those from all-atom simulations (Figs. 2b–c, f–g and S11b–c, g–h). For highly flexible loops, such as CDR H3 of the anti-HBeAg e6 Fv, the dynamics remain limited compared to those sampled using enhanced sampling techniques like metadynamics as these methods can explore conformational states separated by high energy barriers, regions that a residue-level model like CV3-Fv may not fully access. We also observed that the frame-wise RMSD distributions from the CV3-Fv simulations lacked the multimodal characteristics occasionally seen in all-atom simulation profiles, displaying instead more Gaussian-like profiles. This suggests that CV3-Fv simulations may not accurately reproduce either the balance between distinct conformational states or the energy barriers that separate them, limiting the accuracy of the predicted kinetics. However, this behaviour is not surprising, given that coarse-grained models are known to smoother the protein energy landscape (Kmiecik et al., 2016; Jin et al., 2022), often due to a lack of convergence.

**Figure 2.**
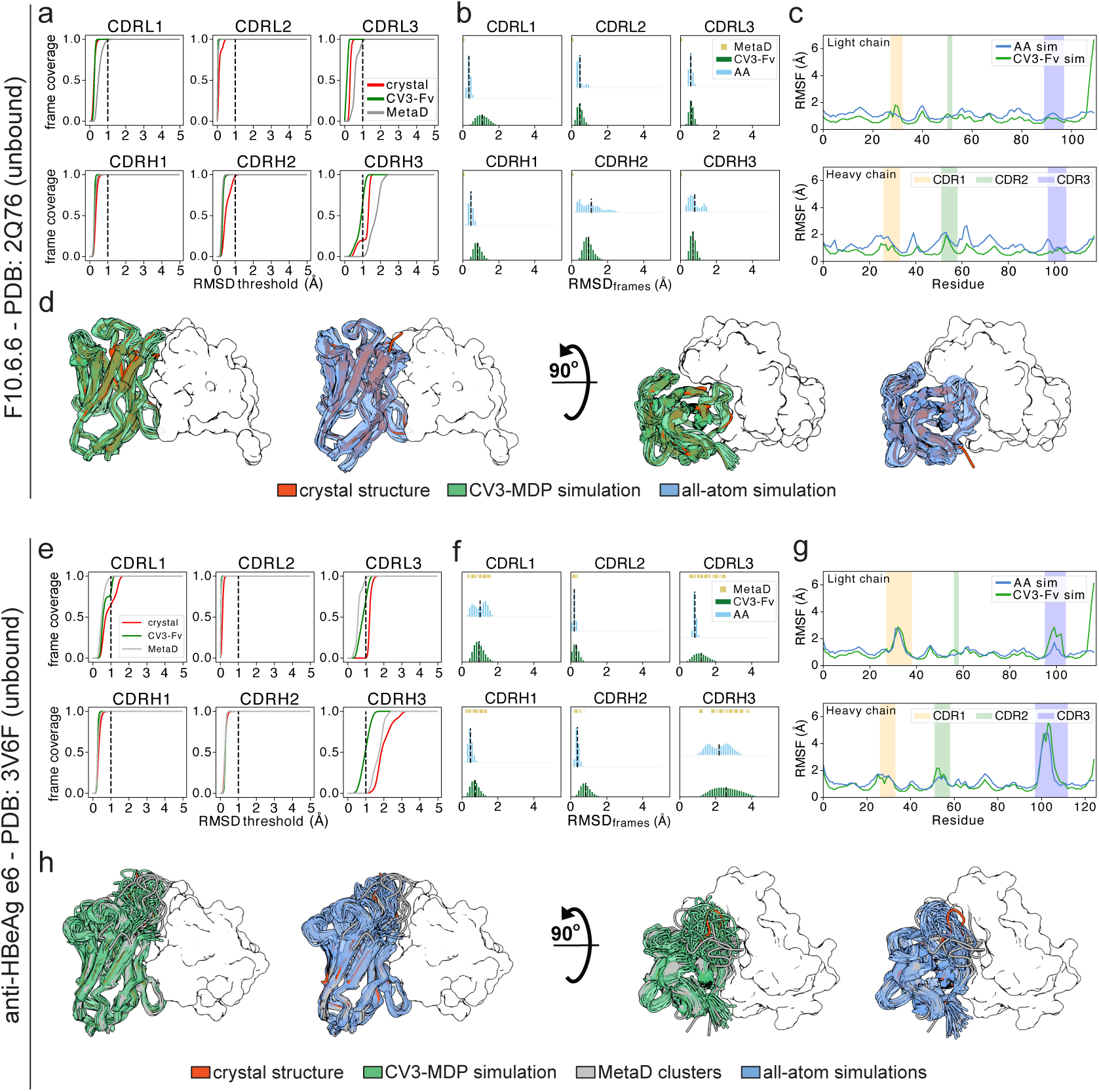
Benchmarking the accuracy of CV3-Fv simulations against all-atom and metadynamics simulations. Comparison of the dynamical properties of Fv regions, particularly the CDR loops, for two monoclonal antibodies: anti-hen egg white lysozyme (PDB: 2Q76) and anti-HBV e-antigen (PDB: 2Q76). (a,e) ‘Frame coverage’ showing the proportion of all atom frames covered at a given threhold by any CV3-Fv frame (green) as well as for the metaD cluster structures (blue) and the starting crystal structure (red). The fraction of all-atom covered frames is reported on the y-axis and the corresponding threshold distance on the x-axis. A black dashed line indicates the 1Å threshold. (b, f) Framewise root-mean-square deviation (RMSD) distribution for all six CDR loops, comparing CV3-Fv (green), all-atom simulations (light blue), and metaD cluster structures (yellow). Mean RMSD values are indicated with black dashed lines. (c, g) Per-residue RMSF values from CV3-Fv (green) and all-atom (blue) simulations. Shaded backgrounds mark different CDR loops. (d, h) Structural ensembles of the Fv region: light chains are shown as white surfaces and heavy chains as cartoons. For the heavy chain, CV3-Fv (green) and all-atom (blue) representative structures are overlaid the top of the starting crystal structure (red) and metaD clusters. Left and right panels show side and top views, respectively.

We further extended our CV3-Fv benchmark by analysing simulation for four additional Fvs: three from antigen-bound crystal structures and one from an unbound Fv. Weighted cluster structures from metaD simulation were available for each system for comparison. We found good agreement between our CV3-Fv simulations and the metaD structures in terms of predicted flexibility for each CDR (Figs. S5 and S4). However, we observed that, in all simulations (including those of our first two benchmark systems), the CDRH2 loops exhibited greater conformational diversity in our simulations than in the metaD clusters, suggesting a possible overestimation of its flexibility. Nevertheless, given the short length outside the framework of CDRH2 (Kelow et al., 2022) and its limited functional role, the impact of this overestimation is expected to be minimal.

### Comparison with ensembles observed in experimental structures

As multiple experimental structures are available for several antibodies, often capturing distinct conformational states, we examined how well the CDR conformational space, flexibility and the distribution of VH-VL orientation angles observed in our CV3-Fv simulations match the experimental data.

We investigated the covered conformational space. We extracted all Fvs with multiple conformations in any of the CDRs from SAbDAb (numbers for each CDR are shown in Fig. 3a) and assessed their coverage in CV3-Fv simulation using ‘alternative conformation coverage’. This metric measures the minimum distance from any simulation frame to each experimental conformation except for the conformation from which the simulation was started (Fig. 3a). We compared against a baseline of the same metric computed from only the starting structure and found that our simulations consistently outperformed this measure. Specifically, CV3-Fv simulations sampled over 80% of experimentally observed CDRH1 and L1 conformations and over 90% of H2, L2 and L3 conformations below 1Å. For CDRH3 coverage was slightly lower, with coverage of around 60% of conformations below 1Å.

**Figure 3.**
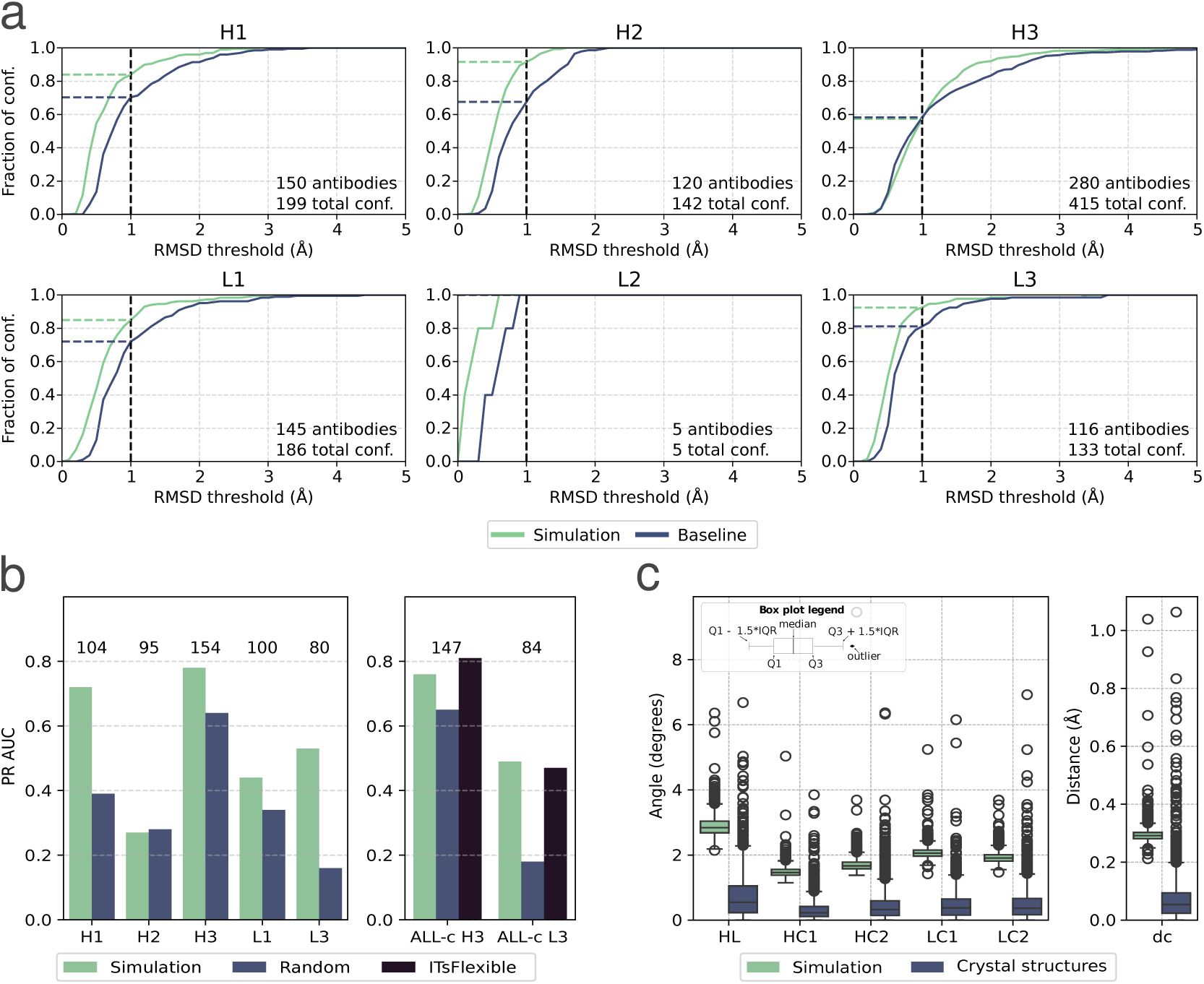
Comparison of simulations against experimental structural ensembles. (a) Alternative conformation coverage of CV3-Fv simulations. The plot shows the percentage of alternative conformational states (all conformation excluding the conformation of the starting structure) covered by CV3-Fv at each distance threshold (green). The baseline (blue) provides a comparison by calculating the same metric based on only the starting structure. (b) Prediction of CDR flexibility from RMSD observed in CV3-Fv simulations. PR AUC for classification of rigid and flexible CDRs (determined in crystal ensembles) from frame pairwise RMSD in CV3-Fv simulations (green). Analysis was performed the for five CDRs calculating RMSDs using the standard IMGT CDR definitions (H1, H2, H3, L1, L3). Performance was also analyzed on the ALL-conformations CDRH3 and CDRL3 datasets which uses a modified CDR definition. The Classification is compared to a random baseline (blue) and the ITsFlexible method (black). The number of samples is reported above each bar. (c) Distribution of VH-VL orientation angles (HL, HC1, HC2, LC1, LC2) and inter domain distance (dc). The distribution of the metrics in CV3-Fv simulations (green, 1764 samples plotted) and crystal structure ensembles (blue, 3140 samples plotted) was calculated and the standard deviation is shown.

To compare the CDR flexibility in the simulations to experimental data, we used a previously described approach (Spoendlin et al., 2025). CDRs of Fv structures from SAbDAb with multiple available structures were assigned binary labels (Spoendlin et al., 2025). Each CDR was labelled as either “rigid” or “flexible”, depending on whether a conformational change of more than 1.25 Å was observed between different crystal structures (see Methods for details). This approach does not guarantee that all CDRs labelled “rigid” do not exhibit conformational flexibility as it is possible that some conformational states have not been captured. However, the set should be heavily enriched for CDRs that lack flexibility. We analysed if the magnitude of the frame-pairwise RMSD in the simulation agrees with these labels. Specifically, we used RMSD as an input of a logistic regression and monitored the area under the precision-recall curve (PR AUC; Fig. 3b). As no CDRL2s were labelled as “flexible” we only performed analysis for the remaining five CDRs (the number of datapoints are indicated in the figure). PR AUC predicted from CV3-Fv RMSDs exceeds a random baseline for four of the five CDR. For CDRH2, we observed a value close to random, which may be explained by our previous observation that the CV3-Fv protocol overestimates CDRH2 flexibility. We further analysed performance on ALL-conformations (Spoendlin et al., 2025) which allows for comparison against ITsFlexible (Spoendlin et al., 2025). ITsFlexible is a deep learning tool specifically trained for this binary classification task and was benchmarked on the ALL-conformations data set. ALL-conformations contains CDRH3 and CDRL3 loops assigned “rigid” and “flexible” labels as in the data sets introduced here with a small difference in the IMGT residues over which RMSD is calculated. On both datasets, CV3-Fv achieves results comparable to ITsFlexible. Taken together, these results indicate a good agreement of flexibility in CV3-Fv simulations and experimental data.

Lastly, we compared the distribution of VH-VL orientation angles (HL, HC1, HC2, LC1, LC2) and the inter domain distance (dc) to observations in crystal structure ensembles. Our simulations tend to sample a slightly wider distribution of angles (1-2 degrees) and distances (0.3 Å) than observed in the experimental ensembles (Fig. 3c). In absolute terms these are small deviations and overall the method captures the ensemble well.

### Investigating the limitation of our flexibility predictions

Monoclonal antibodies can undergo a range of conformational changes upon binding to their specific antigens. The extent and nature of these changes depend on several factors, including affinity maturation and the structural constraints imposed by the antigen (Zhao et al., 2019). Moreover, for the purpose of crystallization, antibodies are often bound to small molecules, peptides or partial fragments of the antigen. As a result, paratopes may exhibit significantly different conformations compared to the unbound state, with smaller buried surface areas and fewer hydrogen bonds compared to antibody–antigen complexes (Lee et al., 2022). Therefore, we assessed how the choice of starting structure effects simulation using three Fvs. For each of these, crystal structures representing different starting conditions (unbound, bound, bound in alternative conformation) were available.

First, we examined Efalizumab Fv, which we found in three diffeent conformations in the PDB: two from a bound state (PDB ID: 3EOA) and one from an unbound state (PDB ID: 3EO9). The CDRs in these structures were moderately similar, with RMSDs consistently below 1 Å for all CDRs. We found consistent flexibility predictions for all CDRs across all three simulations, regardless of the initial structure (Fig. S8) including CDRH3 (Fig. S8c). When we compared our results with cluster structures from metaD simulations, we found that the CV3-Fv simulations depict CDR H3 as a more rigid loop than the metaD conformations suggest. It is interesting to note that, despite the metaD clusters sampling a broader and more flexible ensemble, they do not include conformations close to the initial unbound crystal structures (Fig. S8c,e).

As small conformational changes between starting structures did not impact the outcome of simulations, we continued our investigation with Fvs showing larger differences (RMSD > 1 Å) in CDR conformation. We used Fvs of antibodies D3h44 and AL-57 as two example systems.

For D3h44 Fv, we collected four distinct conformations: one from an unbound structure (PDB ID: 1JPT) and three from structures bound to two different antigens – another Fv (PDB ID: 1PG7) and a tissue factor (PDB ID: 1JPS). The conformation for the CDRH3 loops differ between the bound and unbound state, but also between the two complexes, with RMSD over 1 Å. We found good agreement between flexibility metrics across simulation starting from the different crystal conformations (Fig. S6a,b). As in previous cases, we also found that CV3-Fv predicted flexibility of CDRH3 was underestimated compared to the metaD conformation. However, when we inspected the conformational ensembles, in particular for CDRH3, we observed significant differences between the ensembles (Fig. S6c,d), with ensembles generated from the unbound structure and the Fv-Fv complex differing significantly from those generated from the complex with the tissue factor. We observed the same pattern in the metaD simulations, which were initiated from the tissue factor–complex crystal structure and did not sample conformations close to the unbound state.

We saw similar patterns in the second system analysed, the Fv of AL-57, which had three available crystal structures — two in complex with an antigen (PDB: 3HI6) and one unbound (PDB 3HI5). Here, the difference in CDRH3 conformations between the bound and unbound structures is over 1.5 Å. As before, we observed minimal differences in the flexibility metrics obtained for the different CV3-Fv simulations (Fig. S7a, b), regardless of the initial structure. However, when we examined the structural ensemble of CDRH3 (Fig. S7c,d,e), we found that simulations initiated from the bound state did not sample the conformational space of the unbound crystal structure. Again, this observation was consistent with the cluster centroids from the metaD simulations. We then examined the torsion angles of the CDRH3 loop and identified a small number of residues with distinct distributions of these angles for specific residues, particularly Asp109 and Phe115 (Fig. S7f), located near the termini of the loop and close to the characteristic CDR “kink” turn (Weitzner et al., 2015). Here, simulations initiated from the bound conformation explored a torsional space markedly different from that of the unbound state and were unable to access it. The inability of the bound simulations, even in the structures from metaD simulations, to reach the unbound state suggests that these torsion angles may be separated by high energy barriers. This, combined with differences in the harmonic restraints typically applied to CDR residues (which depend on the starting crystal conformation), may further restrict their ability to explore alternative conformations.

Overall, these results suggest that starting from a bound or unbound conformation might lead to different conformational bundles, even when flexibility metrics remain consistent across simulations. This is particularly evident when bound structures are used as starting points, as CDR loops may undergo substantial conformational changes that place them in regions of the energy landscape separated from the unbound state by significant energy barriers.

### FlAbDab and FTCRDab

Having established that our CV3-Fv protocol captures the flexibility of CDR loops, we created the Flexibility Antibody Database (FlAbDab) and the Flexibility TCR Database (FTCRDab) two large data-bases of simulated antibodies and TCRs. We initially simulated all non-redundant and quality filtered experimentally solved structures before a cutoff data (May 29th 2024 for antibodies, April 10th 2024 for TCRs). This led to simulations of 3140 antibody Fvs (including standard Fvs and single chain Fvs) and 280 TCR Fvs. We further increased the size of FlAbDab by simulating an additional dataset of 148,000 antibody structures predicted with ABodyBuilder2 and released as part of their original publication (Abanades et al., 2023). The CV3-Fv protocol runs with a low computational cost of approximately 2 hours on a standard CPU for a single antibody or TCR. This allowed us to create the two databases using approximately 300,000 hours of CPU computation.

### Analysis of CDR dynamics

Although the primary focus of this work was the generation of a new dataset we also carried out a brief characterisation of the CDR flexibility from the simulations of antibodies and TCRs started from experimentally determined structures. This set contained 3140 antibody and 280 TCR Fvs.

We used RMSD and RMSF metrics to characterize the dynamics of antibody CDRs. Specifically, we used the RMSD to the mean simulation structure (RMSD_mean_) and residue-level RMSF. As expected the highest RMSD_mean_ is observed for CDRH3 with a median of 0.9 Å (Figure 4), as it tends to be the longest CDR.. However, even for this loop the majority of samples show only moderate flexibility with only few outliers exhibiting RMSD values of above 2 Å. CDRH2 shows the second highest RMSD_mean_, which may seem surprising as H2s tend to be shorter than most other CDRs. This finding may be linked to CV3-Fv tendency to overestimate H2 flexibility. However, nearly all residues defined as CDR2 by IMGT are actually loop residues. By contrast, IMGT-defined CDR1 and CDR3 loops frequently include residues located in the β-strands at either end of the loop. This structural difference is likely to be responsible for the relatively high RMSD values observed for CDR2. CDRs H1, L1 and L3 show a similar magnitude of flexibilities with mean RMSD_mean_ of around 0.6 Å. A very low RMSD_mean_ is observed for CDRL2, which is explained by an average CDRL2 length of 3 residues in our dataset. We observed a strong correlation between the loop RMSD_mean_ and CDR length (Pearson R 0.77) as well as more specifically for CDRH3 RMSD_mean_ and length (Pearson R 0.77).

**Figure 4.**
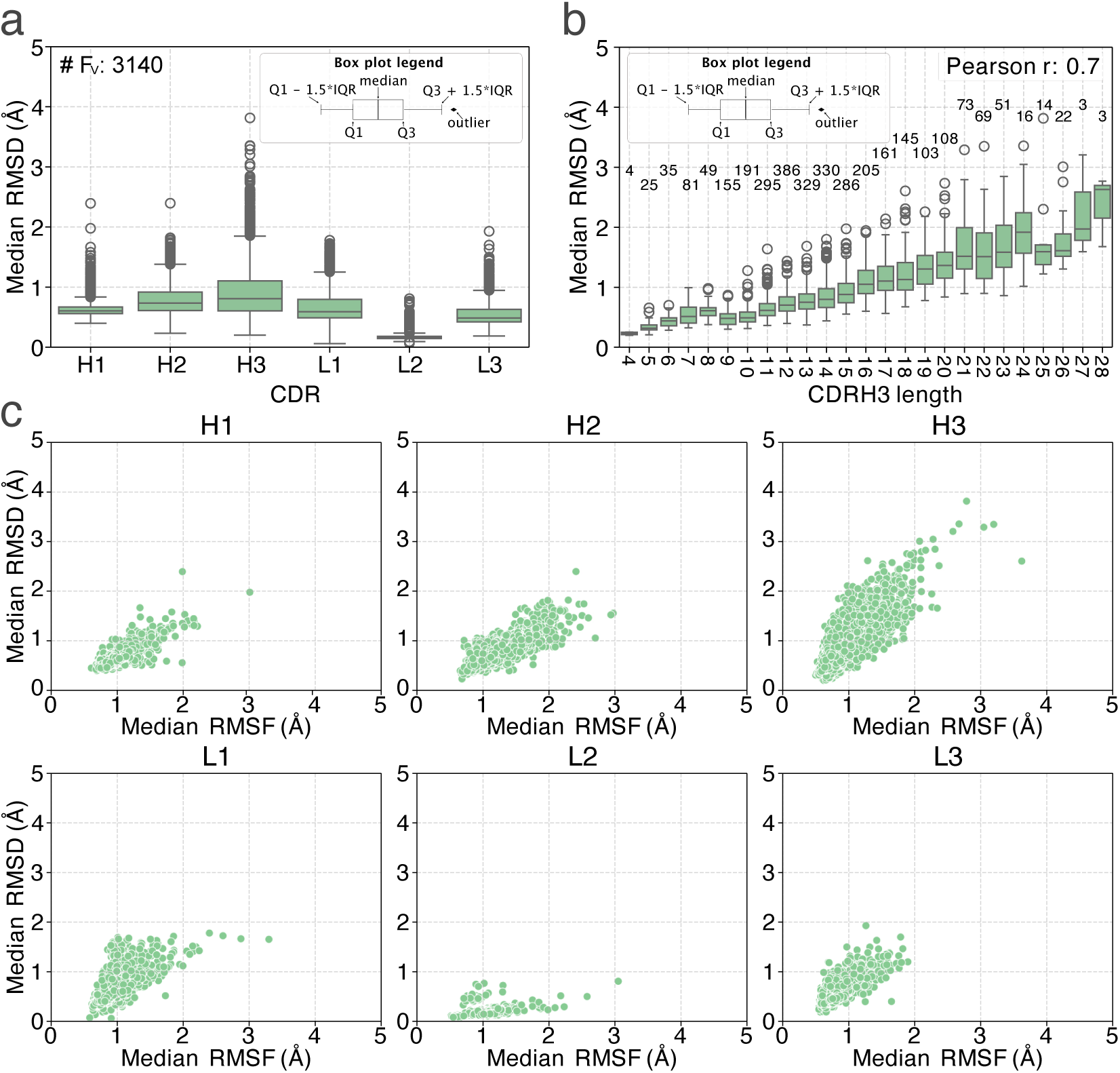
CDR dynamics of antibody with experimental structures. CV3-Fv simulations were run for 3140 non-redundant and quality filtered antibody structures from the SAbDab. a) Box plot showing the median RMSD across all frames to the the average loop structure (RMSD_mean_) for each of the CDRs. Each box plot shows values of all 3140 samples. b) Box plot showing the median CDRH3 RMSD_mean_ as a function of CDR length. A Pearson correlation of RMSD against CDRH3 length was run and the correlation coefficient indicated. The number of samples is reported on the top of each box plot. c) Scatter plot showing the median RMSD_mean_ of each loop against the median residue-level RMSF.

We further analyzed the relationship between CDR RMSD_mean_ and residue-level RMSF and saw two regimes of loop movements. In the first, we observed high RMSD and low RMSF values (Figure 4c) which indicates a large structural change with little fluctuation over time, representing slow transitions between dissimilar conformational states. This regime appears most frequently in CDRH3. For the other five CDRs, we more commonly observed a second regime described by low RMSD and high RMSF values which describes faster frequency oscillations around a more narrow conformational state.

We then investigated whether our data shows a connection between antibody maturity and flexibility. Several studies (*Schmidt et al., 2013*; *Babor and Kortemme, 2009*; *Li et al., 2015*; *James et al., 2003*; *Blackler et al., 2022*; *Fernández-Quintero et al., 2019b*) have suggested CDR rigidification upon maturation while others argue against it (*Jeliazkov et al., 2018*). First, we confirmed that our simulations (Fig. S9) had successfully characterised the rigidification of CDR loops previously reported in specific antibody maturation lineages (*Fernández-Quintero et al., 2020*). We then investigated if antibodies more distant from the germline show lower flexibility. We could not establish correlation between RMSD_mean_ and the sequence identity to the V and J genes of the corresponding germline for any of the CDRs (Figure S10a & b). As lower sequence identity does not strictly mean an antibody is more mature, we also examined whether germline antibodies in general are more flexible than matured antibodies. We labelled an antibody chain as ‘naive’ if both V and J genes were identical to the germline and ‘mature’ if either gene contained more than 3 mutations. Comparing RMSD_mean_ of mature and naive antibodies, we observed that naive antibodies were on average more flexible than mature ones. However, the differences were small and we could not establish statistical significance. Taken together, our data does not show a clear trend of rigidification with maturation.

We also analysed the TCR simulations. CDRs B3 and A3 show the highest RMSD_mean_ values in our data (Figure S13). These tend to be the longest loops in TCRs and exhibit the highest sequence diversity (Quast et al., 2025). On average, we found slightly lower RMSD values compared to the antibody set. As we again observed a correlation between length and flexibility, this finding can be attributed, at least in part, to the shorter length of TCR CDRs. We also highlight that most TCR CDRs exhibit low RMSD and high RMSF indicating faster oscillations around a narrower conformational state.

## Discussions

Structural dynamics of antibodies, particularly in the Fv region and CDR loops, are thought to be essential for their function. This flexibility is tightly linked to binding affinity (Mikolajek et al., 2022; Li et al., 2015; Schmidt et al., 2013; Haidar et al., 2014) and may enable the structural adaptability required to recognise diverse antigens (James et al., 2003; Fernández-Quintero et al., 2019; Guthmiller et al., 2020). Therefore, flexibility is a key factor in the discovery and design of novel therapeutics. However, the limited availability of data on Fv dynamics makes it difficult to study these effects systematically and even more challenging to train predictive models to address the millions of uncharacterised antibody sequences.

Recent advances in computational power and coarse-grained modelling have made it possible to use MD simulations to generate large datasets for exploring protein dynamics on a large scale (von Bülow et al., 2025a). In this study, we examined the suitability of CALVADOS 3, a residue-level coarse-grained model initially designed for folded domains and intrinsically disordered regions, for simulating the dynamics of antibody Fv regions.

We found that simulating Fv by considering the CDR loops to be entirely intrinsically disordered regions led to an overestimation of their flexibility in CALVADO3, so we introduced additional simulation restraints within the CDR loops based on the interaction patterns observed in the initial crystal structure. This new setup, called CV3-Fv, generates ensembles that showed greater agreement in terms of flexibility and conformational sampling with all-atom and enhanced sampling simulations. Although flexibility metrics correlate well with the atomistic data, CV3-Fv simulations tend to underestimate distinct conformational states and the barriers between them, particularly in highly flexible loops. Consequently, we sometimes found that the relative populations of macrostates were different for more flexible systems compared to all-atom simulations. This limitation is not surprising, given that coarse-grained models inherently smooth the underlying protein energy landscape (Kmiecik et al., 2016; Jin et al., 2022).

We also investigate the effect of using bound or unbound crystal structures as starting points for the simulations. We found that structures of bound Fv, particularly those of Fvs bound to antigen fragments or peptides, can lead to distinct conformational ensembles, even when flexibility metrics report similar values. This divergence is particularly apparent when the initial structures differ in terms of the backbone torsion angles within the CDRs, which results in different local constraints during the simulation. Consequently, certain regions of the conformational space, especially those near the native unbound state, may not be fully sampled. While this issue could stem from our strategy for introducing constraints into the model, we observed similar behavior in metadynamics simulations, which also failed to sample the unbound region when initiated from a bound crystal structure. Recent studies (Zhao et al., 2019; Lee et al., 2022) have shown that some CDR loops can undergo substantial conformational changes upon antigen binding. These shifts may result in states separated by high energy barriers from the unbound conformation, which makes it difficult, or even impossible, for simulations such as CV3-Fv to recover the unbound ensemble. While not all complexes exhibit this behaviour, it is important to note that over 90% of antibody structures in SAbDab are in the bound form. This suggests that this limitation may affect a significant proportion of the simulated Fvs.

We used CV3-Fv to created FlAbDab, the first database of molecular dynamic simulations that covers all non-redundant, experimentally resolved antibody Fvs (over 3000 simulations) in addition to a two orders of magnitude larger set of predicted structures (150,000 simulations). We expanded our work to create FTCRDab which comprises MD simulation for all non-redundant, experimentally resolved TCR structures (280 simulations). The release of FlAbDab and FTCRDab opens the possibility for several downstream applications. The databases will allow the study of CDR dynamics and its functional importance in a large set of immune receptors and provide a new data source for training machine learning models.

We performed a preliminary analysis on the subset of antibodies and TCRs with experimentally resolved structures. For the majority of samples, we observed moderate magnitudes of flexibility with a median RMSD_mean_ below 1 Å for all CDRs and only a few outliers showed values higher than 2 Å. A correlation between loop length and flexibility highlights length as a key factor in determining loop dynamics, although previous studies suggest that length alone is not sufficient to explain flexibility (Spoendlin et al., 2025; Guloglu and Deane, 2023). Our large datasets may allow the identification of sequence or structural motifs with the potential to more rationally engineer CDR flexibility (Kouba et al., 2024).

While the functional importance of CDR flexibility has been investigated in case studies (*Miko- lajek et al., 2022*; *Li et al., 2015*; *Schmidt et al., 2013*; *Haidar et al., 2014*; *James et al., 2003*; *Fernández-Quintero et al., 2019*; *Guthmiller et al., 2020*), the extent to which flexibility is exploited in natural immune repertoires to balance between affinity and specificity remains poorly understood. It has been suggested that naive antibodies tend to be more flexible to promote promiscuous, low-affinity binding and thereby increasing the effective size of the immune repertoire and that during maturation antibodies become more conformationally restricted (*Schmidt et al., 2013*; *Babor and Kortemme, 2009*; *Li et al., 2015*; *James et al., 2003*; *Blackler et al., 2022*; *Fernández-Quintero et al., 2019b*). In contrast, a previous large-scale study has not found any evidence that maturation promotes rigidity (*Jeliazkov et al., 2018*). Similarly, although our simulations reproduced the phenomenon of rigidification in lineages known to exhibit it, we found no clear, universal trend of increased CDR rigidity when comparing naïve and mature antibodies. However, we highlight several caveats in our analysis. Firstly, analysis was performed on SAbDab antibodies which do not present a natural immune repertoire and patterns of flexibility may differ. Secondly, our analysis was limited to statistics across a large number of simulations and, as shown in previous studies, rigidification may well be relevant to individual antibody lineages in a more nuanced manner. The investigation of the functional role of CDR flexibility at repertoire level using our databases of antibody MD simulations will be an important future direction.

Another application of our databases is its use for training machine learning model. Despite the recent developments (Lewis et al., 2025; Zhu et al., 2024; Zheng et al., 2024; Lu et al., 2023; Huguet et al., 2024; Jing et al., 2024; Janson et al., 2025), accurately predicting protein conformational states remains an open challenge. Although few models have been developed to predict dynamics independently of MD data (Cagiada et al., 2025; Invernizzi et al., 2025), most existing approaches follow a similar principle: they employ generative structure prediction frameworks that include at least one training phase based on MD-generated ensemble structures. However, despite promising results indicating that conformational dynamics can be learned, the amount of available MD data appears to be at least one of the limiting factors. The largest previously available MD simulation databases are one to two orders of magnitude smaller than the number of solved protein structures and do not contain antibodies or TCRs (Vander Meersche et al., 2024; Mirarchi et al., 2024; Tesei et al., 2024). Our databases provide more than 150,000 simulated immune receptors and promises the potential of improving conformation prediction for these proteins. While models trained on our datasets may inherit the same biases observed in the CV3-Fv simulations, our database is a valuable pretraining resource. One possible future approach to addressing these biases could be to first train models using FlAbDab trajectories, and then fine-tune them using either higher-accuracy MD simulations or limited experimental data.

Together, FlAbDab and FTCRDab offer new possibilities for the systematic exploration of the sequence-structure relationship in antibodies and TCRs, as well as the principles behind adaptive immunity, and also facilitate the development of next-generation computational models of immune recognition.

## Methods

### CV3-Fv workflow

For each query Ab and TCR, the FV structures to use as input for the simulation (VH and VL domains) were extracted from the PDB file and the remainder of the structure, including conserved domains and antigens, was discarded. Then, we run using OpenMM a short energy minimization of Fv structure (Eastman et al., 2024) with a protocol identical to the one used as part of AF2 (Jumper et al., 2021). The protocol used AMBER ff99SB force field and additional harmonic constraints with a spring constant of 0.7 N/m were placed on all non-hydrogen atoms. Minimization was performed until the system reaches a tolerance of 2.39 kcal/mol.

We performed molecular dynamics simulations in the NVT ensemble at 25 °C and pH 7 using the CALVADOS suite (von Bülow et al., 2025b). Simulations used a Langevin integrator with a time step of 10 fs and a friction coefficient of 0.01^-1^ ps. Salt-screened electrostatic interactions were modelled using a dielectric constant of 74.2 and an ionic strength of 0.15 M. Simulations were conducted in a periodic box with dimensions of 25 × 25 × 150 nm for a total of 100.5 ns. The 100 ns length was chosen as a compromise to achieve convergence in trajectories and computational efficiency (Fig. S14)

In the first instance, we used the residue model CALVADOS 3 using parameters from the official release (Cao et al., 2024), following the functional forms described in the original CALVADOS publication. Here, regions outside the IMGT-defined CDR loops were treated as folded domain and restrained using a harmonic potential with a force constant (k) of 700 N/m. We named this setup CALVADOS 3-IMGT (CV3-IMGT). Given, the poor agreement with all-atom simulation in predicting the flexibility of CDR region, we introduced additional harmonic restraints to selected residue pairs within loops under two conditions: (i) if a hydrogen bond was present between the atoms of the pair in the crystal structure or (ii) if their side-chain centres of mass were in close proximity. Restraints were applied using a harmonic potential with a force constant of k = 175 N/m for the former and k = 350 N/m for the latter. Hydrogen bonds and side-chain contacts within CDRs were identified using the DSSP algorithm. A hydrogen bond was defined as present if the DSSP energy score was lower than −0.41 kJ/mol for a given acceptor-donor pair. Side-chain interactions were defined as any residue pair with a centre-of-mass distance below 4.5 Å (using the Cα atom for glycine). We named this setup CALVADOS 3-Fv (CV3-Fv).

Coarse-grained simulations were converted to all-atom trajectories using the cg2all package (Heo and Feig, 2024) with the ‘ResidueBased’ option selected to convert the residues from beads to all atoms. The representative ensemble for each entry in the FlAbDab and FTCRDab databases includes frames randomly selected from the simulation trajectories. The total number of frames in each ensemble is proportional to the flexibiluty of CDRH3, ranging from a minimum of 5 structures for rigid entries (RMSD < 0.5 Å) to a maximum of 15 structures for highly flexible entries (RMSD > 3 Å).

### Other datasets

#### All atom MD simulations

The all atom simulations of TCR PDB 7f5k (chains A and B) were obtained from a previous study (Quast et al., 2025) where a single replicate of a 2 μs unrestraint simulation was performed at 300K and pH 7.5. We perfomed simulations for antibodies anti-hen egg white lysozyme (PDB: 2Q76) and anti-HBV e-antigen (PDB: 2Q76) as well as Titin Epitope in HLA-A1 (PDB 5bs0) as outline below.

We performed complete all-atom MD simulations of mouse anti-hen egg white lysozyme antibody F10.6.6 Fab fragment (PDB: 2Q76), monoclonal anti-HBV e-antigen Fab fragment (e6), unbound (PDB: 3V6F), and the Titin Epitope in HLA-A1 (PDB: 5BS0). All non-protein residues were removed from the file. The original anti-HBV e-antigen Fab fragment PDB file contained insertion codes in chain H and L, which was renumbered to follow the sequential IMGT numbering scheme, i.e., assigning consecutive residue numbers based on the IMGT antibody numbering convention. The original F10.6.6 Fab fragment PDB file contained missing electron densities in chain B residues 130–134 which was modelled using Chimera X. The structures were protonated at the pH they were crystallised at using H++ (Anandakrishnan et al., 2012), and the AMBER compatible file was extracted from this.

We ran all simulations with AMBER24 (Case et al., 2023) using the ff19sb force field (Tian et al., 2019) with the OPC3 water model (Izadi and Onufriev, 2016), and all systems were prepared using tleap from the AmberTools package (Case et al., 2023). For each structure,we manually defined disulphide bonds between the appropriate cysteine residues to maintain a proper tertiary structure. We prepared the solved systems through a multistep process in which the protein charge was first neutralised, followed by solvation in a truncated octahedral OPC water box with a buffer distance of 10.0 Å from the protein surface to the edges of the box. We finally added additional randomised sodium and chloride ions to achieve a physiological ionic strength of approximately 150 mM, with ion concentrations adjusted based on the initial protein charge state.

All solvated systems underwent multistage relaxation and thermal equilibrium prior to production MD. We perfomed initial energy minimisation for 1000 cycles using a hybrid approach with 30 cycles of steepest descent followed by conjugate gradient optimisation. We applied strong harmonic positional restraints of 100 kcal mol^−1^ Å^−2^ to all protein atoms while allowing freedom for solvent molecules and ions to optimise their positions around the constrained protein structure. We used a 10 Å cutoff for nonbonded interactions during this initial minimisation phase.

We then heated the systems from 100 K to 298 K over 1 ns (1,000,000 steps with 1 fs time steps) using canonical ensemble (NVT) MD with Langevin temperature regulation. We used Langevin thermostat used a collision frequency of 1.0 ps^−1^ to provide efficient temperature coupling while maintaining the same 100 kcal mol^−1^ Å^−2^ restraints on protein atoms. Temperature ramping was implemented using the nmropt facility with linear scaling from the initial temperature of 100 K to the target temperature of 298 K over the entire 1 ns duration.

Following thermal equilibration, we performed isothermal-isobaric (NPT) equilibration on the system at 298 K and 1 atm pressure for 1 ns using the Monte Carlo barostat algorithm with the same protein restraints maintained. We additionally performed an additional 1 ns NPT equilibration with reduced protein restraints of 10 kcal mol^−1^ Å^−2^ to allow initial structural relaxation while maintaining overall protein integrity.

We carried out a second phase of energy minimisation consisting of 10,000 cycles with selective application of restrictions only to the protein backbone atoms (C_α_, N and C atoms) at 10 kcal mol^−1^ Å^−2^, allowing side chain optimisation while preserving the secondary structure. The nonbonded cut-off was reduced to 8 Å for all subsequent simulation stages.

We then achieved a progressive restraint reduction through a series of 1 ns NPT equilibration runs with backbone restraints that steadily decreased from 10 kcal mol^−1^ Å^−2^ to 1 kcal mol^−1^ Å^−2^ to 0.1 kcal mol^−1^ Å^−2^, allowing gradual increase in protein flexibility and conformational sampling. Each stage used Langevin dynamics for temperature control with a collision frequency of 1.0 ps^−1^ and the Monte Carlo barostat for pressure regulation.

In the final equilibration stage we performed a 1 ns of completely unrestrained NPT MD to achieve complete system relaxation and establish baseline dynamics prior to production simulations. All equilibration simulations employed SHAKE constraints to fix bond lengths involving hydrogen atoms, particle-mesh Ewald summation for long-range electrostatic interactions, and periodic boundary conditions. We saved trajectory coordinates 10 ps for analysis, with system energies and properties monitored every 1 ps to ensure stability throughout the equilibration protocol.

Following equilibration, we ran three replicates for antibody systems and four for TCR systems, performing MD production simulations for 500 ns under NPT conditions at 298 K and 1 atm pressure. The production simulations employed a 2-fs integration timestep with SHAKE constraints applied to all hydrogen-containing bonds to enable a larger timestep while maintaining numerical stability. The temperature regulation was maintained using the Langevin thermostat with a collision frequency of 1.0 ps^−1^, and pressure control was achieved through the Berendsen weak coupling algorithm. The cutoff for non-bonded interactions was extended to 9 Å for production simulations to improve the accuracy of long-range interactions. Trajectory coordinates were saved every 10 ps (5000 steps) for subsequent analysis, with system energies and thermodynamic properties recorded at the same frequency. We generated checkpoint files every 100 ps to enable simulation continuation and recovery. We did not apply any positional restrictions during production simulations, allowing complete conformational freedom for all components of the system, including protein, solvent, and ions.

#### Metadynamics simulations

We used a dataset of 9 antibodies (PDBs: 2q76, 3v6f, 3hi6, 3eoa, 1jps, 5i18, 5i15, 3g6d, 2y07) simulated with all-atom metadynamics simulations released by (Spoendlin et al., 2025) for evaluation. Simulations details are reported here from the published protocol (Fernández-Quintero et al., 2019b). In a first step, well-tempered metadynamics was used for enhanced sampling. A linear combination of the sine and cosine ψ torsion angles of CDR loops were defined as collective variables. In a second step, representative structures from the metadynamics simulations were used as starting points for short classical MD simulations to obtain unbiased trajectories. Classical MD simulations were clustered based on CDR RMSD with an average linkage hierarchical clustering algorithm and RMSD cut off of 2.5 Å to extract the representative structures we used for evaluation.

#### Experimental structures

We ran simulations for all non-redundant and quality filtered antibody Fvs, scFvs and TCR Fvs with experimentally resolved structures. We extracted all antibody and scFv structures from the Structural Antibody Database (SAbDab) on May 29th 2024 and TCR structures from the Structural TCR Database (STCRDab) on April 10th 2024. We filtered structures looking for those with a resolution lower than 3.5 Å and no missing residues between IMGT numbered positions 5 and 128. In case of multiple available structures, we selected one structure for each Fv sequence. We preferred unbound structures and those solved by X-ray crystallography. This resulted in a set of 3033 antibodies, 132 scFvs and 281 TCRs. 3009 antibodies, 131 scFvs and 280 TCRs were successfully simulated.

#### Predicted structural models

In addition to the experimental structures we simulated a larger dataset of predicted structural models. We used the dataset released by (Abanades et al., 2023) of 148,000 non-redundant paired antibody sequences from the OAS (Olsen et al., 2022) predicted with ABodyBuilder2 (Abanades et al., 2023).

### Benchmarking

#### Evaluation against all atom and metadynamics simulation

We used four metrics to compare the CV3-Fv protocol against all-atom and metadynamics simulations: frame-pairwise RMSD, RMSD_mean_, residue-level RMSF and frame coverage. Frame-pairwise RMSD measures the pairwise RMSD between all frames in a simulation. For each CDR, we indipendently aligned simulation frames to the mean simulation coordinates using the Cα atoms of CDR residues. We then computed the full (number of frames x number of frames) pairwise Cα RMSD matrix for each CDR.

RMSD_mean_ evaluates the distance of each frames to the mean structure in the simulation. First, we aligned the simulation frames to the mean Cα coordinates across a simulation. We performed separate alignments and evaluations for the two chains (H and L for the antibodies and alpha and beta for the TCRs). We then evaluated the RMSD of each CDR to the mean coordinates for each frame, using the Cα atoms of each CDR residue.

Residue-level RMSF measures the displacement Cα atoms across the simulation. We aligned the simulation frames to the mean Cα coordinates across a simulation. We performed separate alignments and evaluations for the two chains (H and L for the antibodies and alpha and beta for the TCRs). The standard formula was used to calculate RMSF of all Cα atoms of Fv residues.

Frame coverage measures coverage of the conformational space explored in all atom simulation by the CV3-Fv simulations. We compute the pairwise distance matrix of all frames in the all atom simulations to all frames in the CV3-Fv simulations. We used Cα RMSD of CDR residues after alignment on Cα atoms of CDR residues. We defined the coverage of all atom frames as the minimum distance between an all atom frame and any CV3-Fv frame. and we analysed the fraction of all atom frames covered below a given distance threshold.

#### Evaluation against experimental ensembles

We evaluated simulations against structural ensembles observed in crystal structures. First, we collected antibody and scFv structures in the experimental structures dataset and then we filtered for structures solved by X-ray crystallography. We then grouped the filtered structures by Fv sequence identity.

For each Fv sequence, we calculated the number of observed conformations for each of the six CDRs individually. We used pairwise Cα RMSD of CDR residues after alignment on Cα atoms of CDR residues. We defined a conformation cluster by agglomerative clustering with a complete linkage criterion. We used RMSD thresholds of 0.5 Å and 1.25 Å as a more narrow and a wider definition of a conformation. This selected clustering algorithm means that the RMSD of any two structures within a cluster is below the threshold.

For each CDR, we selected all antibodies with more than one experimental conformation (0.5 Å threshold) and calculate a metric we called alternative conformation coverage. Alternative conformation coverage evaluates how well the CV3-Fv simulations cover the experimentally observed conformational space by measuring the minimum distance between an experimental conformation and our simulations. Specifically, for each cluster (conformation), we calculated the pairwise Cα RMSD of CDR residues after alignment on Cα atoms of CDR residues from all cluster members to all simulation frames. We then selected the minimum distance for each member and defined the mean across all members as a conformation coverage metric. We reported the coverage of all conformations except the one containing the structure from which the simulations was started as our metric of alternative conformation coverage. Lastly, we analysed the fraction of alterative conformations covered below a given distance threshold.

To compare the flexibility observed in the CV3-Fv simulations against the experimental data, we followed the analysis used in (Spoendlin et al., 2025). Each CDR was assigned a label of ‘flexible’, ‘rigid’ or ‘unknown’. CDRs with more than one conformation (1.25 Å threshold) were labeled as ‘flexible’. CDRs with a single conformation observed in more than 5 separate PDB structures were labeled as ‘rigid’. The remaining antibodies were assigned to the ‘unknown’ group. We then assessed how predictive flexibility measures of CV3-Fv simulations are of flexible and rigid labels. We then calculated the frame-pairwise RMSD (Cα RMSD of loop residues after alignment on Cα atoms of loop residues) of simulations. The 95th percentile of the frame-pairwise RMSD was then used as an input for binary classification of loops as ‘rigid’ or ‘flexible’ with a logistic regression. The 95th percentile rather than the maximum frame-pairwise RMSD was used to avoid inflation of the value by rare outliers. To assess the accuracy, we used the area under the precision-recall curve (PR AUC) as evaluation metric for the classification.

We further compared the changes in VH-VL orientation in our simulations against the distributions observed in experimental structures. VH-VL orientation was characterized by 5 angles (HL, HC1, HC2, HL1, HL2) and a distance (dc) as described in a previous work (Dunbar et al., 2013). We then evaluated the standard deviation of the six metrics within a structural ensembles for all Fvs with multiple available structures.

## Data and code availability

CV3-Fv simulations for experimentally resolved structures from the FlAbDab and FTCRDab databases are available on Zenodo (https://doi.org/10.5281/zenodo.17552276). Ensembles of representative structures for all the antibodies in FlAbDab, which were predicted using ABodyBuilder2, are also available on Zenodo (https://doi.org/10.5281/zenodo.17555649). Due to the size of the dataset, raw MD simulation data for ABodyBuilder2 are currently available upon reasonable request, and will be made publicly available once a suitable repository has been identified. Allatom MD simulations of two antibodies and one TCR, along with the associated metadata, can be found in the following Zenodo repository: https://doi.org/10.5281/zenodo.17525665. The CV3-Fv pipeline, analysis code, and datasets required to reproduce the analysis can be found at: https://github.com/oxpig/CDR_MD_simulations.

## Acknowledgments

We are grateful to Fan Cao and Soren von Bülow, for their support and guidance in the use of CV3-Fv, and to Kresten Lindorff-Larsen for helpful comments on the manuscript prior to publication. The research was supported by a Novo Nordisk Foundation Postdoctoral Fellowship (NNF23OC0082912) awarded to MC, by research funding from the UK Engineering and Physical Sciences Research Council (EPSCR) [Grant number EP/S024093/1], Roche and the Royal Commission for the Exhibition of 1851 awarded to FS and KI.

## Supplementary Material

### Supplementary Figures

**Figure S1.**
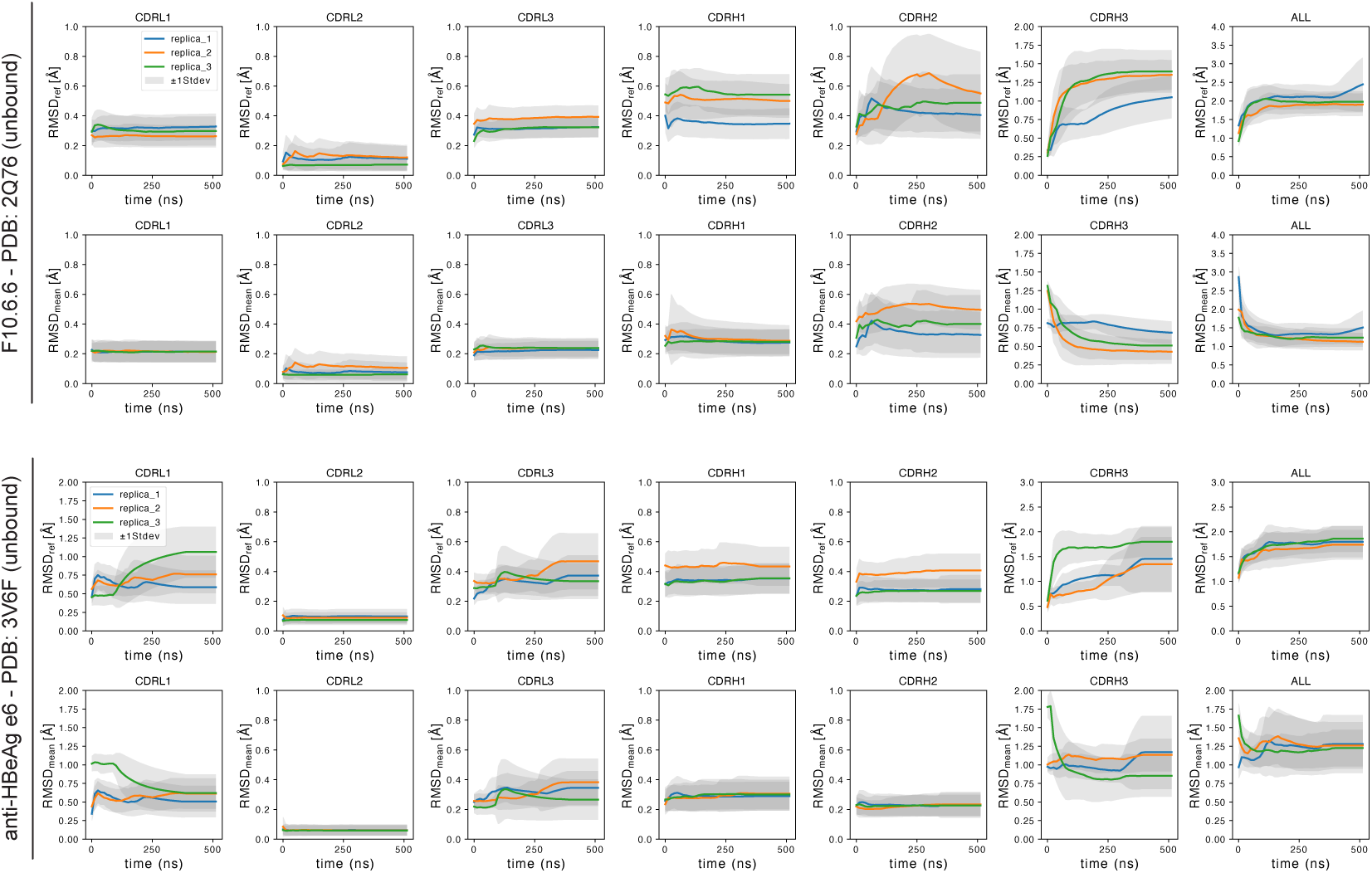
Convergence curves for all-atom simulations of benchmark antibodies. For each antibody system, the upper panels show the cumulative mean RMSD relative to the starting structure for each CDR, while the lower panels show the cumulative mean RMSD relative to the mean atomic coordinates. Each simulation replica is represented by a different colour, and shaded areas indicate the standard deviation around the cumulative mean.

**Figure S2.**
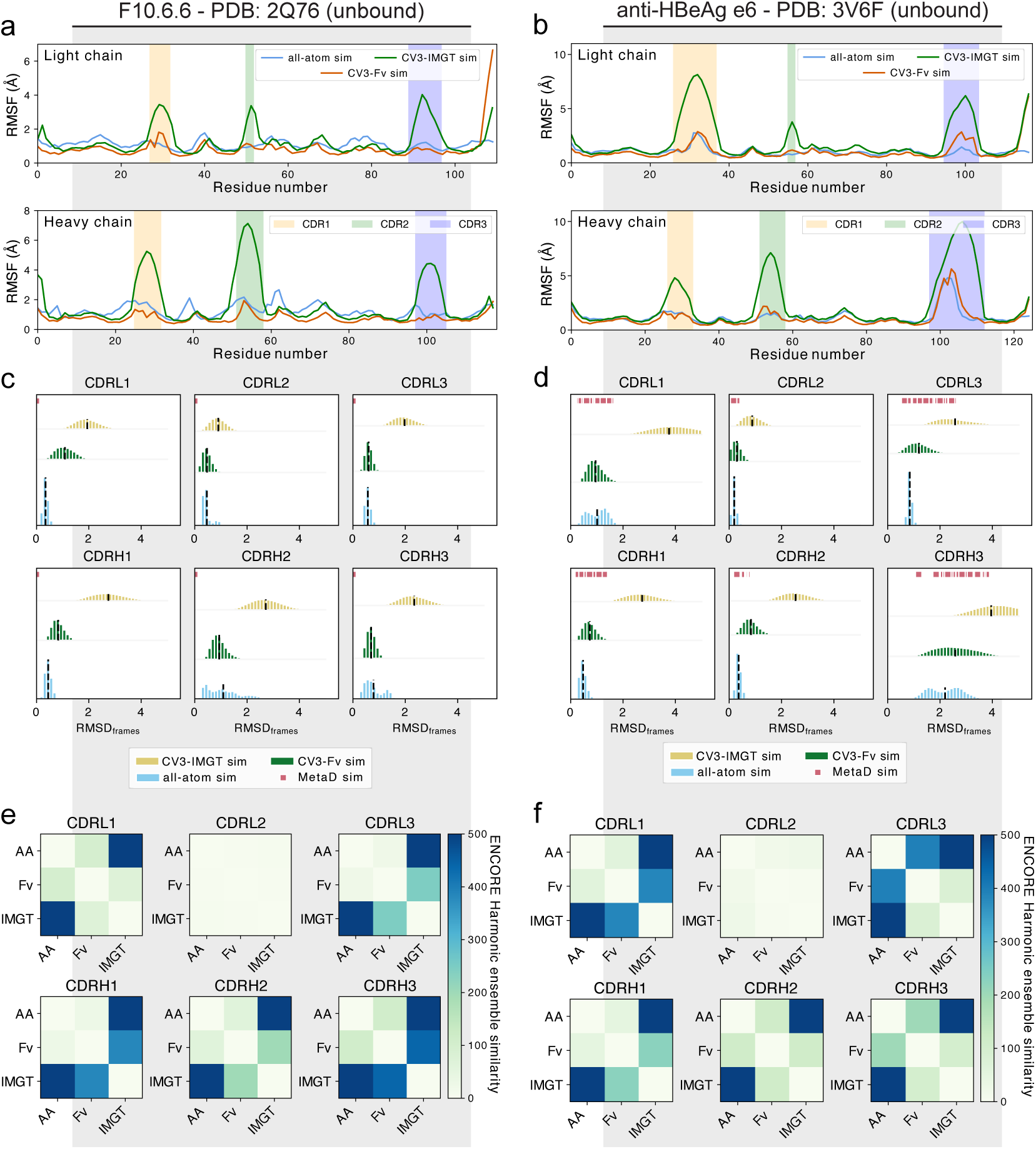
Optimising CALVADOS 3 for Fv simulations Comparison of the flexibility of Fv regions, particularly the CDR loops, in CV3-IMGT and CV3-Fv simulations for two monoclonal antibodies: anti-hen egg white lysozyme (PDB: 2Q76) and anti-HBV e-antigen (PDB: 2Q76). (a,b) Residue-level RMSF values from CV3-IMGT (dark green), CV3-Fv (red) and all-atom (blue) simulations. Shaded backgrounds mark different CDR loops. (c,d) Frame-wise RMSD distribution for all six CDR loops, comparing CV3-IMGT (yellow), CV3-Fv (green), all-atom simulations (light blue), and metaD cluster structures (red). Mean RMSD values are indicated with black dashed lines. (e,f) Harmonic Ensemble Similarity (HES) matrices obtained from the ENCORE package for each CDR in the two benchmark systems, comparing all-atom (AA), CV3-IMGT (IMGT) and CV3-Fv (Fv) simulation ensembles. The lower the HES score, the higher the agreement between the two ensembles.

**Figure S3.**
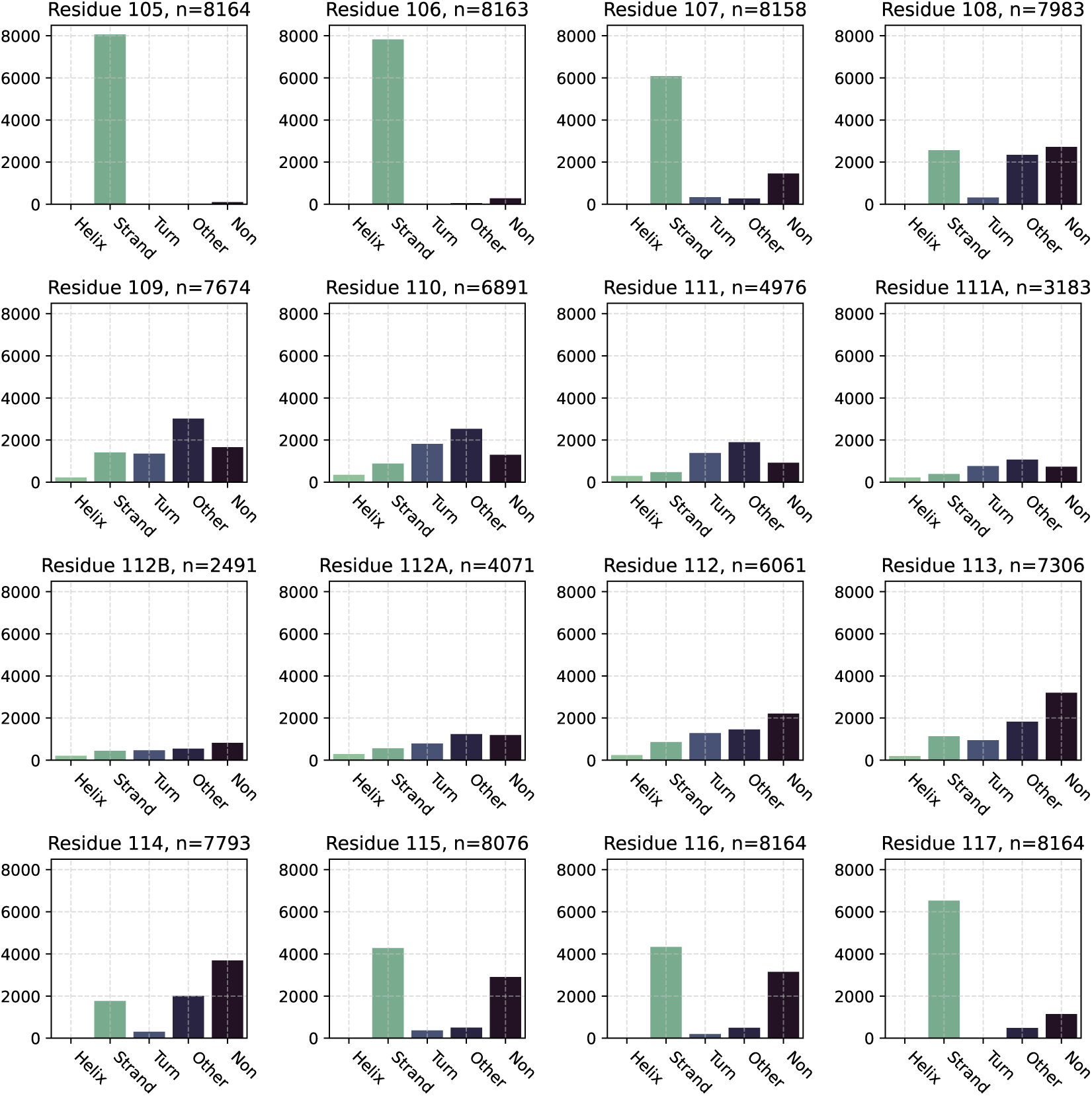
Secondary structure elements in CDRH3 loops The figure shows the type and number of secondary structures present in each IMGT-defined position (one per plot) in CDRH3, as characterised by CV3-Fv in over 3000 experimental structures.

**Figure S4.**
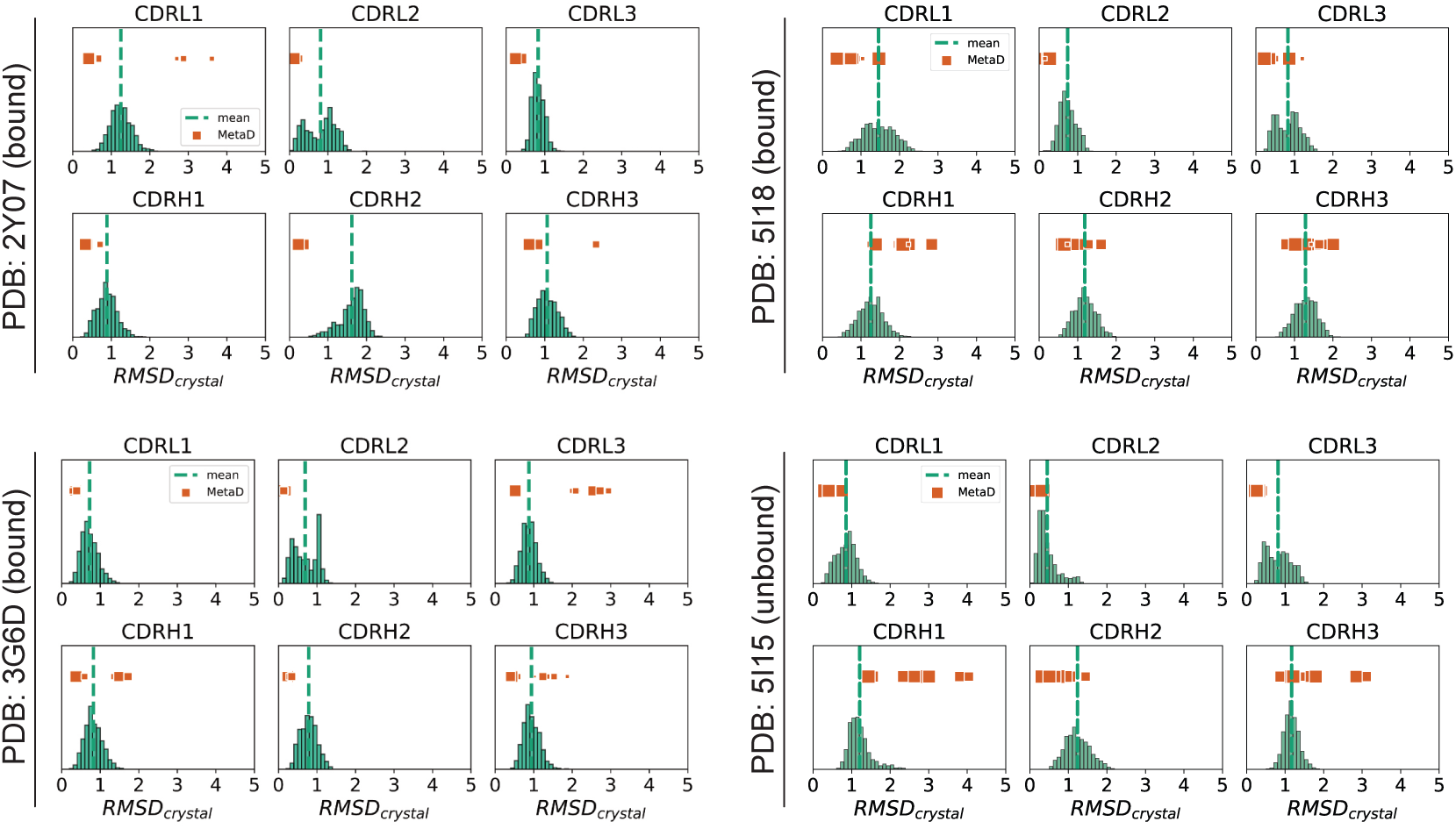
CDR loop RMSD to crystal for CV3-Fv and metaD clusters. The figure shows the flexibility comparison using RMSD from initial structure between CV3-Fv simulations and the metaD cluster structures for four benchmark mAbs, reported here with their crystal structure PDB code. For each system, panels show the RMSD from initial structure distributions for each of the six CDRs. CV3-Fv simulations are shown as green distributions, and metaD cluster values are shown as red dots. The mean RMSD values are shown as green dashed vertical lines in all the plots.

**Figure S5.**
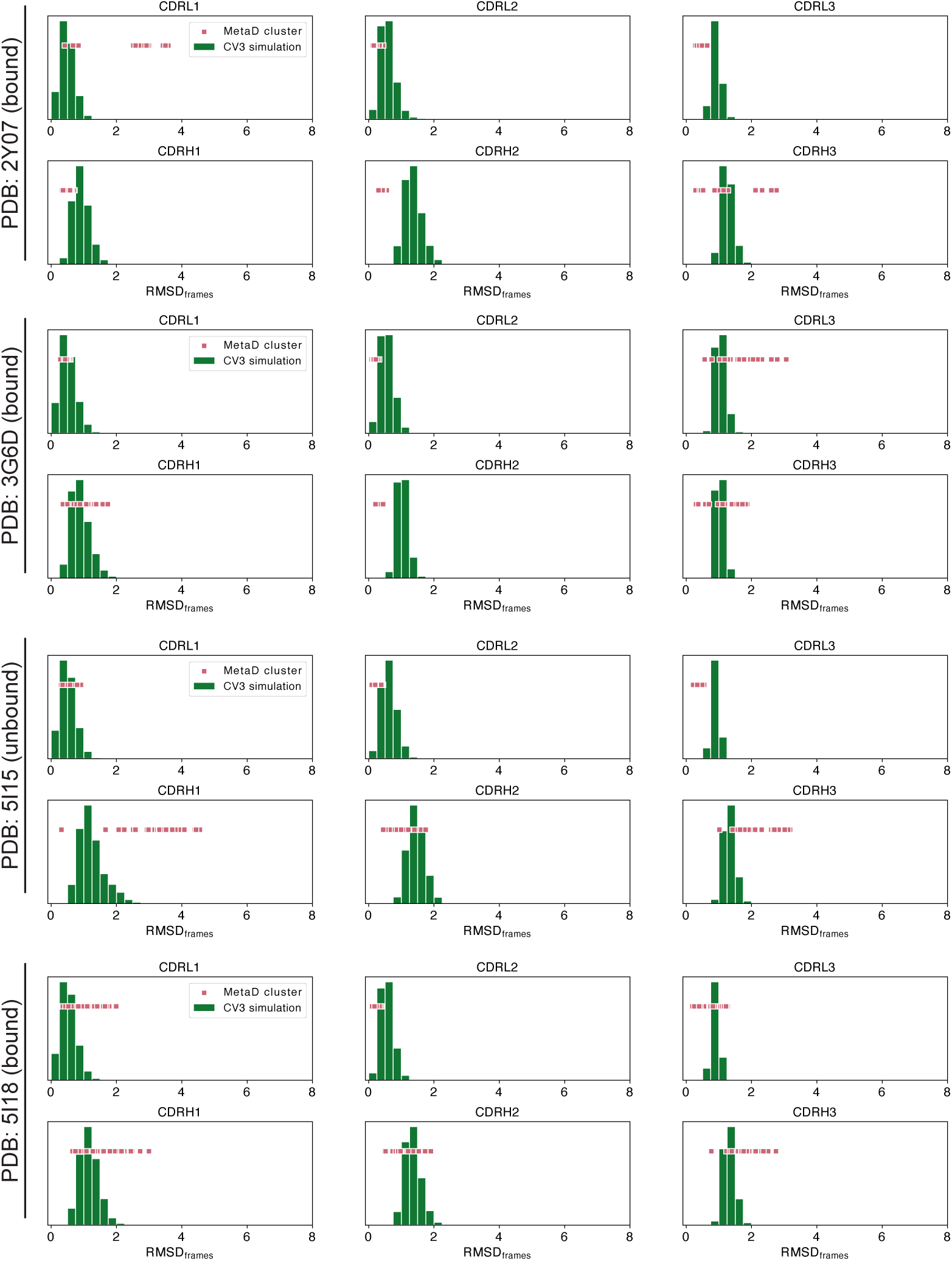
CDR flexibility in CV3-Fv simulations The figure shows the flexibility comparison using RMSD from the mean simulation coordinates between CV3-Fv simulations and the metaD cluster structures for four benchmark mAbs, reported here with their crystal structure PDB code. For each system, each panel shows the RMSD from the mean simulation coordinate distributions for one of the six CDRs. CV3-Fv simulations are shown as green distributions, and metaD cluster values are shown as red dots. The mean RMSD values are shown as green dashed vertical lines in all the plots.

**Figure S6.**
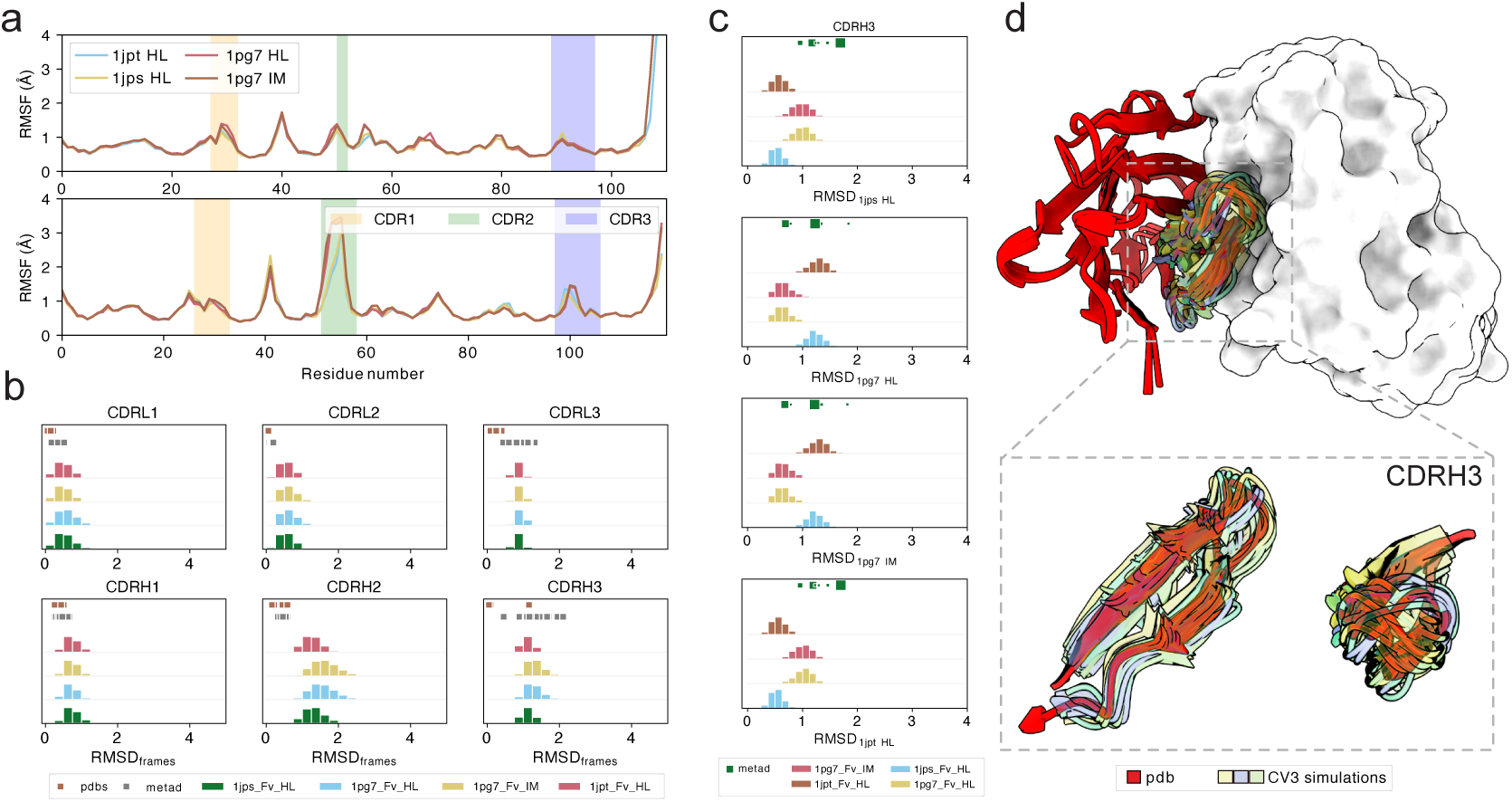
Analysis of the flexibility prediction for the CV3-Fv simulations of the D3h44 Fv from different initial conformations. (a) RMSF values for each residue in the CV3-Fv simulations. Each curve corresponds to a simulation started from a different available crystal structure. Background colours highlight the different CDR loop regions. (b) Frame pairwise root-mean-square deviation (RMSD) distributions for all six CDR loops, comparing CV3-Fv simulations started from different crystal structures (green, yellow, light blue, and red) to metaD cluster structures (gray) and the crystal structures themselves (brown). (c) RMSD distributions from the starting crystal structures for CDRH3, comparing CV3-Fv simulations (coloured as in panel b) to metaD cluster structures (green). (d) Structural illustration of the Fv region. The light chain is shown as a white surface; heavy chains are shown as cartoons. The top panel displays the entire heavy chain from the crystal structure (red) and the CDRH3 loop from CV3-Fv simulations started from different crystal structures (yellow, purple, green). The bottom panel shows a zoomed-in view of the CDRH3 loop.

**Figure S7.**
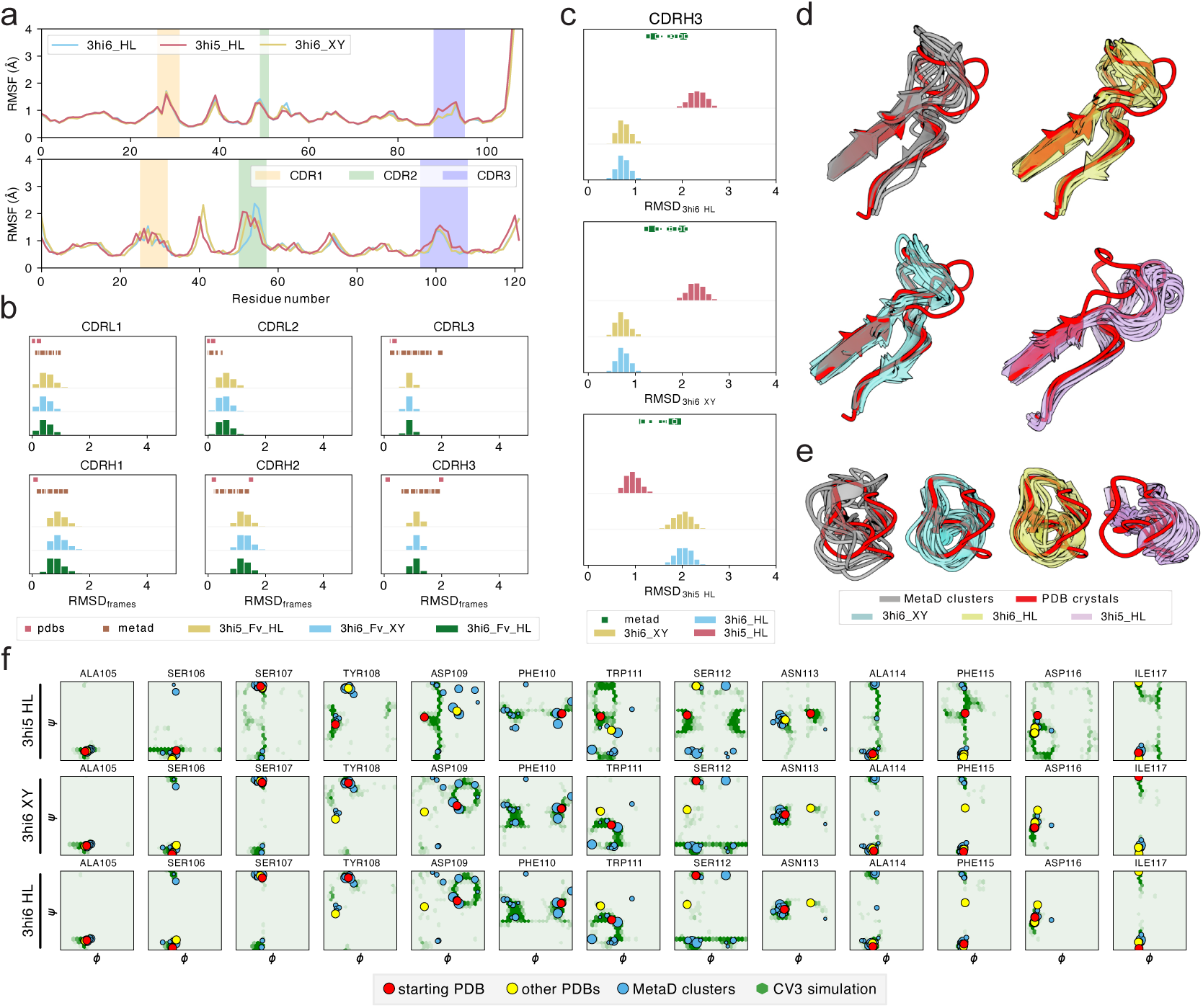
Analysis of the flexibility prediction for the CV3-Fv simulations of the AL-57 Fv from different initial conformations. (a) RMSF values for each residue in the CV3-Fv simulations. Each curve corresponds to a simulation started from a different available crystal structure. Background colours highlight the different CDR loop regions. (b) Frame pairwise root-mean-square deviation (RMSD) distributions for all six CDR loops, comparing CV3-Fv simulations started from different crystal structures (green, yellow, light blue, and red) to metaD cluster structures (brown) and the crystal structures themselves (gray). (c) RMSD distributions from the different starting crystal structures for CDRH3, comparing CV3-Fv simulations initiated from each structure (green, yellow, light blue, and red) to metaD cluster structures (green). (d,e) Structural ensembles of the CDRH3 loop from CV3-Fv simulations and metaD cluster structures. The starting crystal structure is shown in red. Panels (d) and (e) display top and side views of the ensembles, respectively. (f) Torsional backbone space for each residue in CDRH3. Green hex-bin plots show the torsion angles sampled by each residue in our CV3-Fv simulations. Each row corresponds to a different simulation, initiated from a different crystal structure. In each plot, red dots indicate the torsion angles of the starting crystal structure, yellow dots represent alternative crystal structures, and blue dots denote metaD cluster structures.

**Figure S8.**
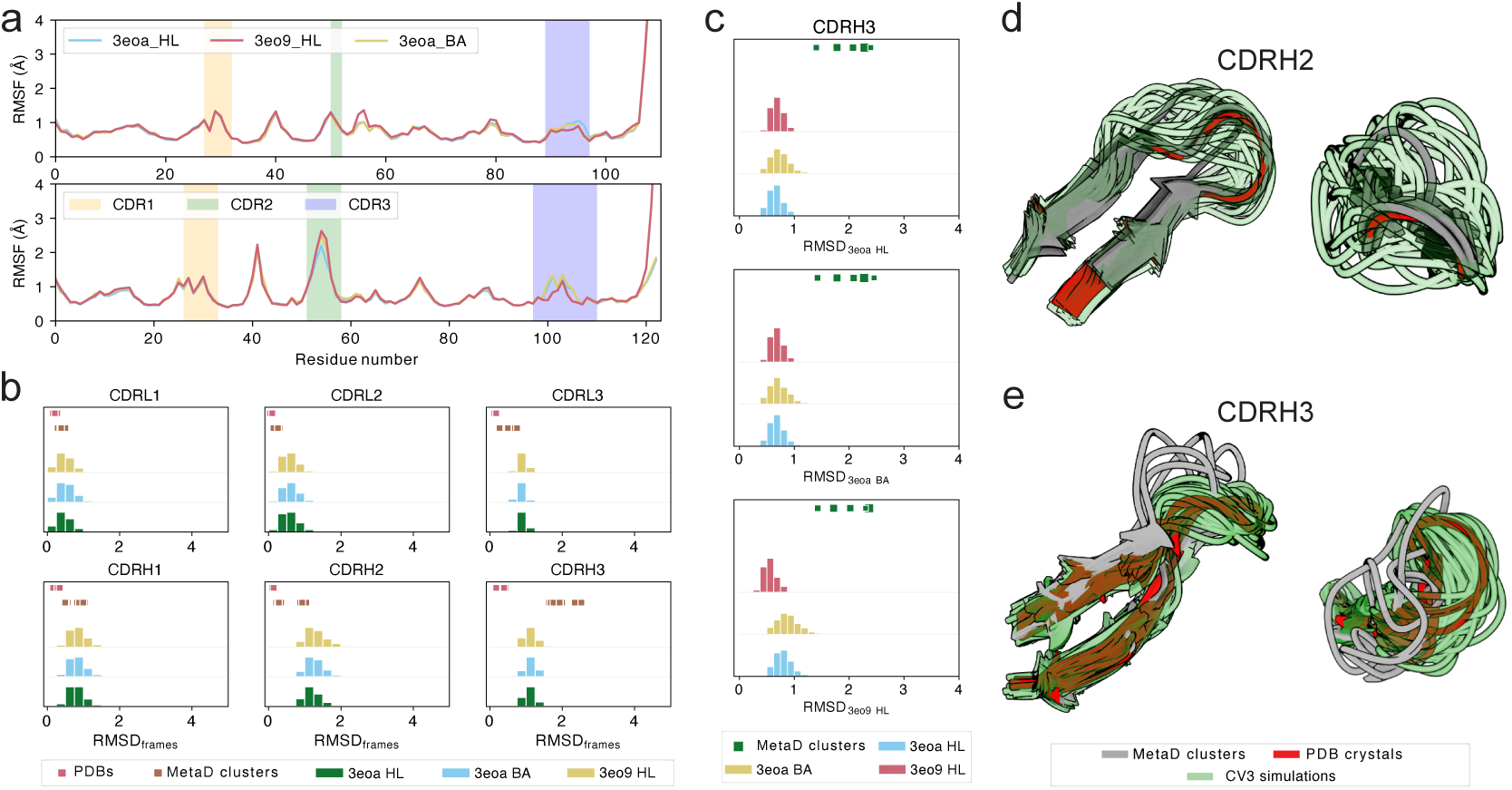
Analysis of the flexibility prediction for the CV3-Fv simulations of the Efalizumab Fv from different initial conformations. (a) RMSF values for each residue in the CV3-Fv simulations. Each curve corresponds to a simulation started from a different available crystal structure. Background colours highlight the different CDR loop regions. (b) Frame pairwise root-mean-square deviation (RMSD) distributions for all six CDR loops, comparing CV3-Fv simulations started from different crystal structures (green, yellow, light blue, and red) to metaD cluster structures (brown) and the crystal structures themselves (red). (c) RMSD distributions from the starting crystal structures for CDRH3, comparing CV3-Fv simulations (coloured as in panel b) to metaD cluster structures (green). (d,e) Representative structural ensembles of the CDRH2 and CDRH3 loops from CV3-Fv simulations (green) and metaD cluster structures. The starting crystal structure is shown in red. Panels on the left and right display top and side views of the ensembles, respectively.

**Figure S9.**
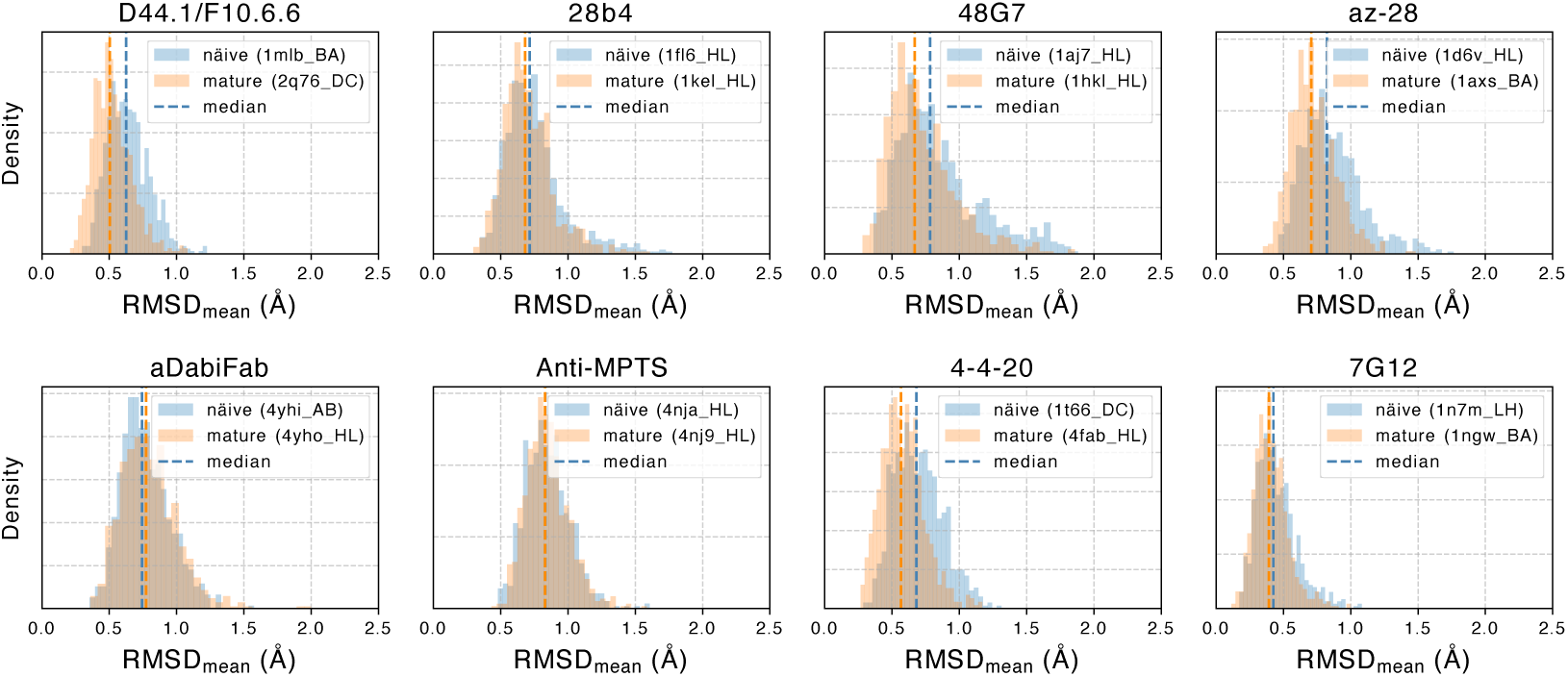
Rigidification of antibody lineages during maturation. Comparison of naive and mature antibodies from the same lineage. Each plot shows the root mean square deviation (RMSD) from the mean coordinates of each frame for the paired naïve (light blue) and mature (orange) antibodies of different lineage types. The vertical lines indicate the median RMSD value.

**Figure S10.**
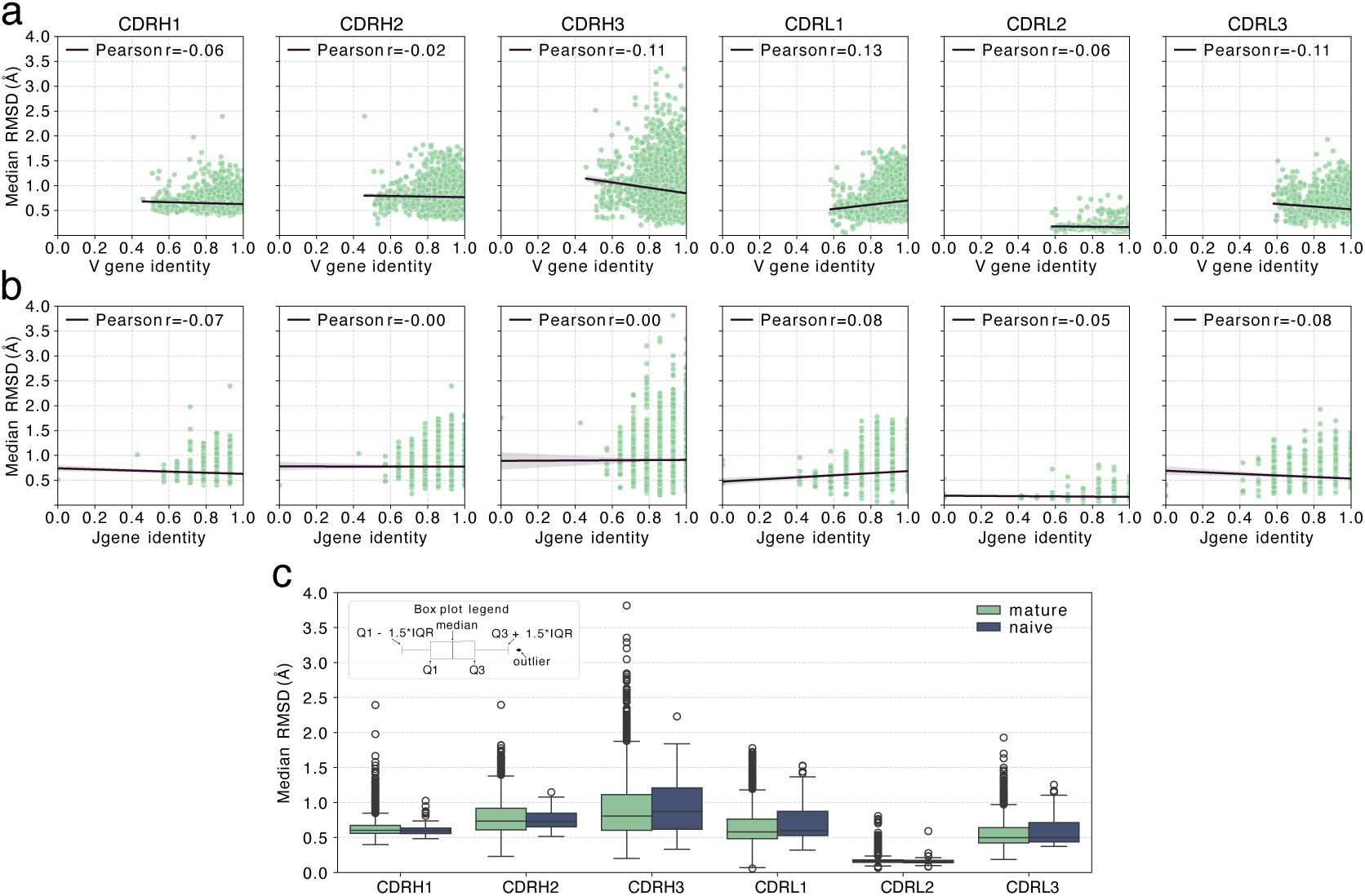
CDR flexibility and antibody maturation. Median CDR RMSD_mean_ plotted against (a) V and (b) J gene identity to the corresponding germline. A linear regression was fit to the data (black line) and the 95% confidence interval is shown in grey shading. Pearson’s correlation coefficient is reported in each panel. c) Distribution of RMSD_mean_ of CDRs in naive (blue) and mature (green) antibody chains. Antibody heavy and light chains were separately classified as naive or mature. The box plot shows samples of 57 naive and 2467 mature heavy chain CDRs (CDRH1-3) and samples of 117 naive and 2020 mature light chain CDRs (CDRL1-L3).

**Figure S11.**
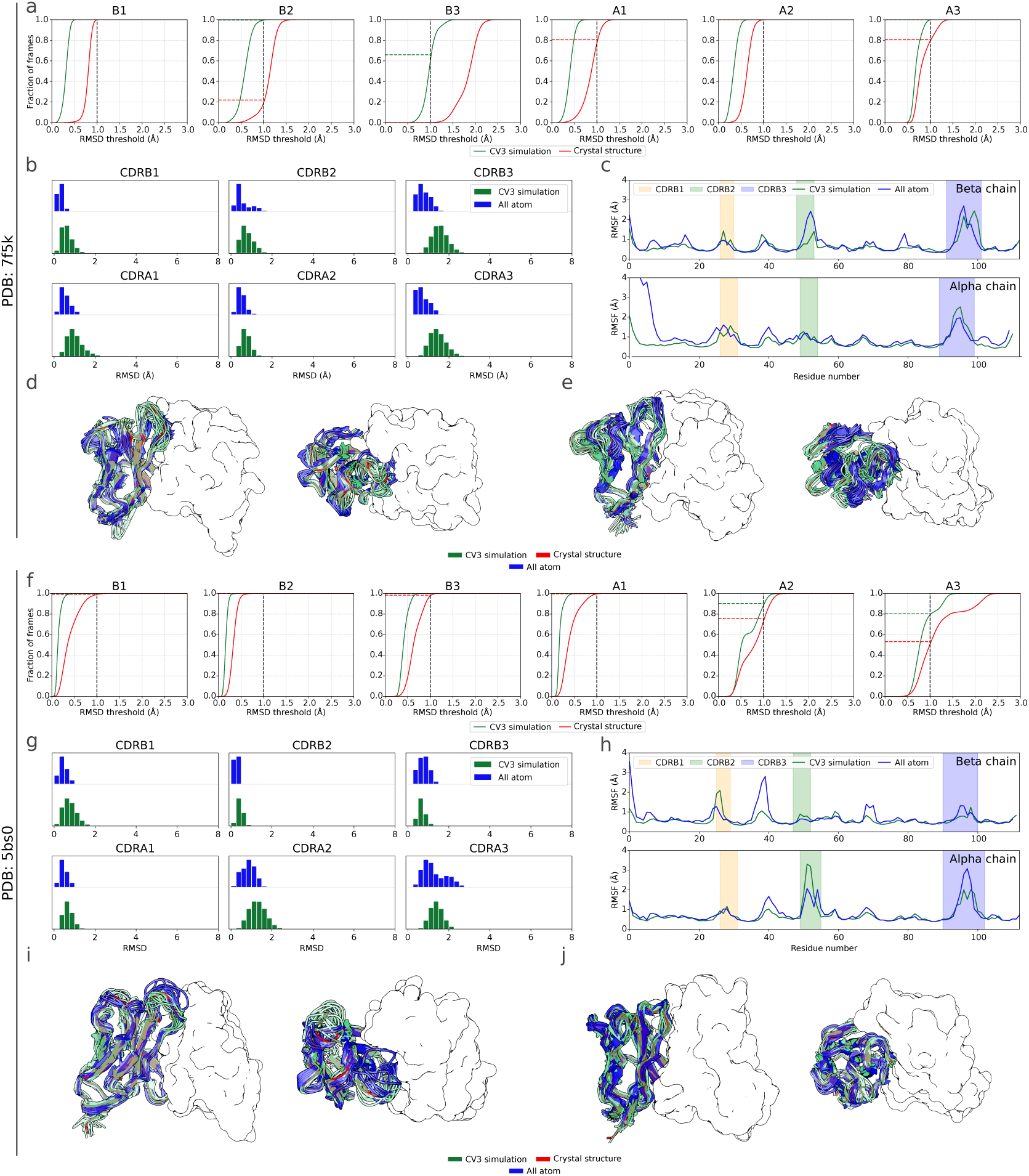
Benchmarking the accuracy of CV3-Fv simulations for TCRs against all-atom simulations. Comparison of the dynamical properties the CDR loops for two TCRs: PDB 75fk and PDB 5bs0. (a,e) Frame coverage showing the proportion of all atom frames covered at a given threshold by any CV3-Fv frame (green) and the starting crystal structure (red). A black dashed line indicates the 1Å threshold. (b, f) Frame pairwise RMSD distribution for all six CDR loops, comparing CV3-Fv (green) and all-atom simulations (blue). (c, g) Per-residue RMSF values from CV3-Fv (green) and all-atom (blue) simulations. Shaded backgrounds mark different CDR loops. (d, i) Structural ensembles of the Fv region. Alpha chains are shown as white surfaces and beta chains as cartoons. For the beta chain CV3-Fv (green) and all-atom (blue) representative structures are overlaid on top of the starting crystal structure (red). Left and right panels show side and top views, respectively. (e, j) Structural ensembles of the Fv region. Beta chains are shown as white surfaces and alpha chains as cartoons. For the alpha chain CV3-Fv (green) and all-atom (blue) representative structures are overlaid on top of the starting crystal structure (red). Left and right panels show side and top views, respectively.

**Figure S12.**
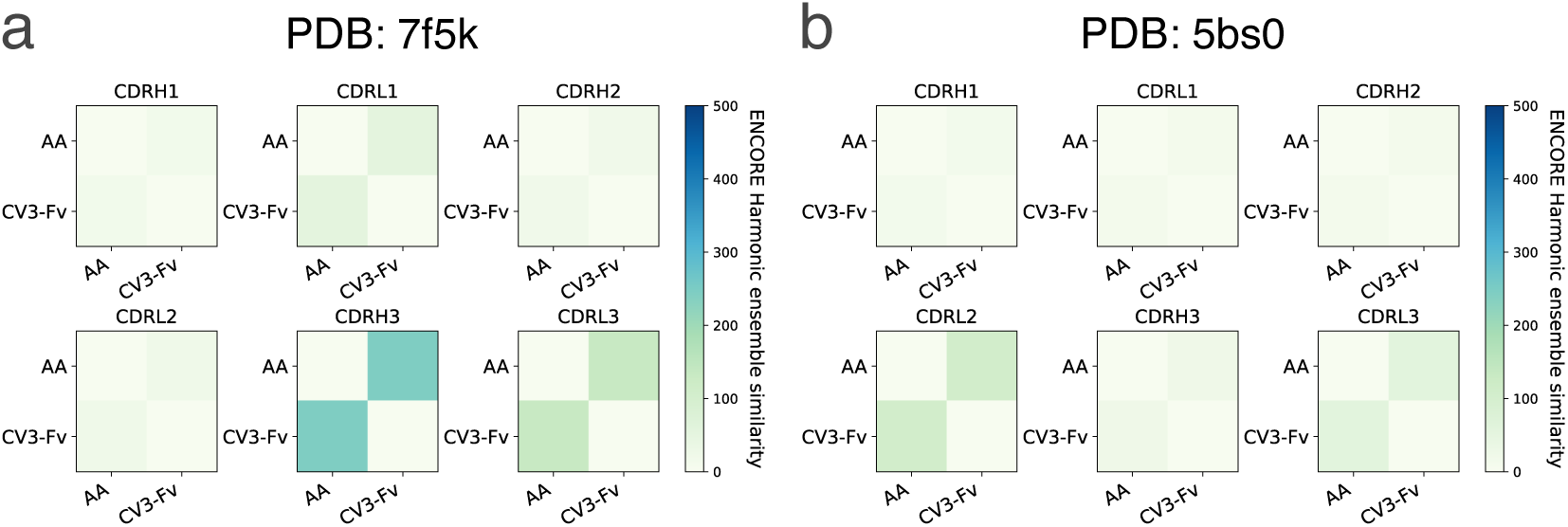
Benchmarking the accuracy of CV3-Fv simulations for TCRs against all-atom simulations with Harmonic Ensemble Similarity (HES). (a,b) HES matrices obtained from the ENCORE package for each CDR in 7f5k (a) and 5bs0 (b), comparing all-atom simulations (AA) and CV3-Fv (Fv). The lower the HES score, the higher the agreement between the two ensembles.

**Figure S13.**
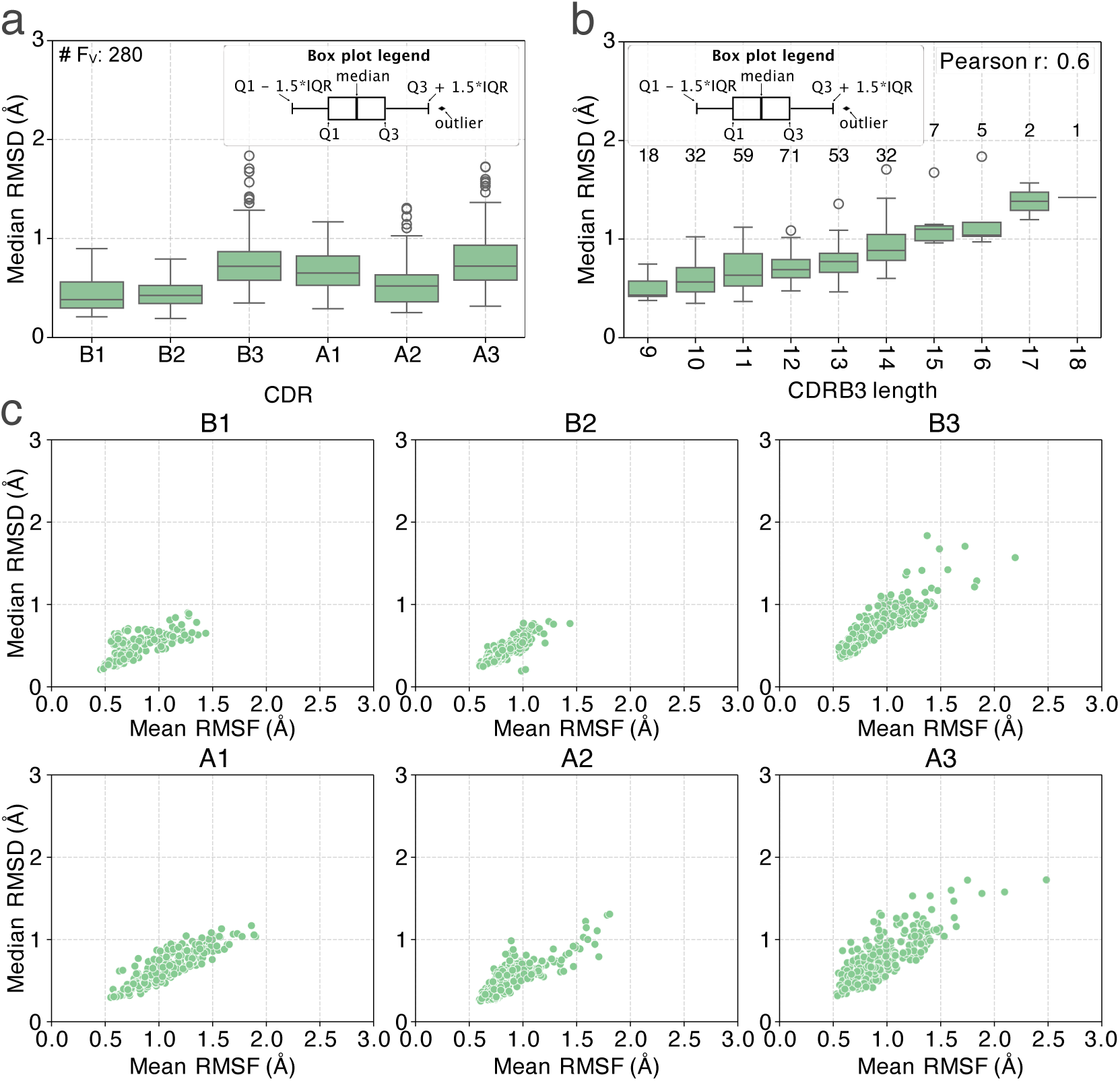
CDR dynamics of TCR with experimental structures. CV3-Fv simulations were run for 280 non-redundant and quality filtered TCR structures from the STCRDab. a) Box plot showing the median RMSD across all frames to the the average loop structure (RMSD_mean_) for each of the CDRs. Each box plot shows values for 280 samples. b) Box plot showing the median CDRH3 RMSD_mean_ as a function of CDR length. A Pearson’s correlation of RMSD against CDRH3 length was run and the correlation coefficient indicated. The number of samples is reported on the top of each box plot. c) Scatter plot showing the median RMSD_mean_ of each loop against the median residue-level RMSF.

**Figure S14.**
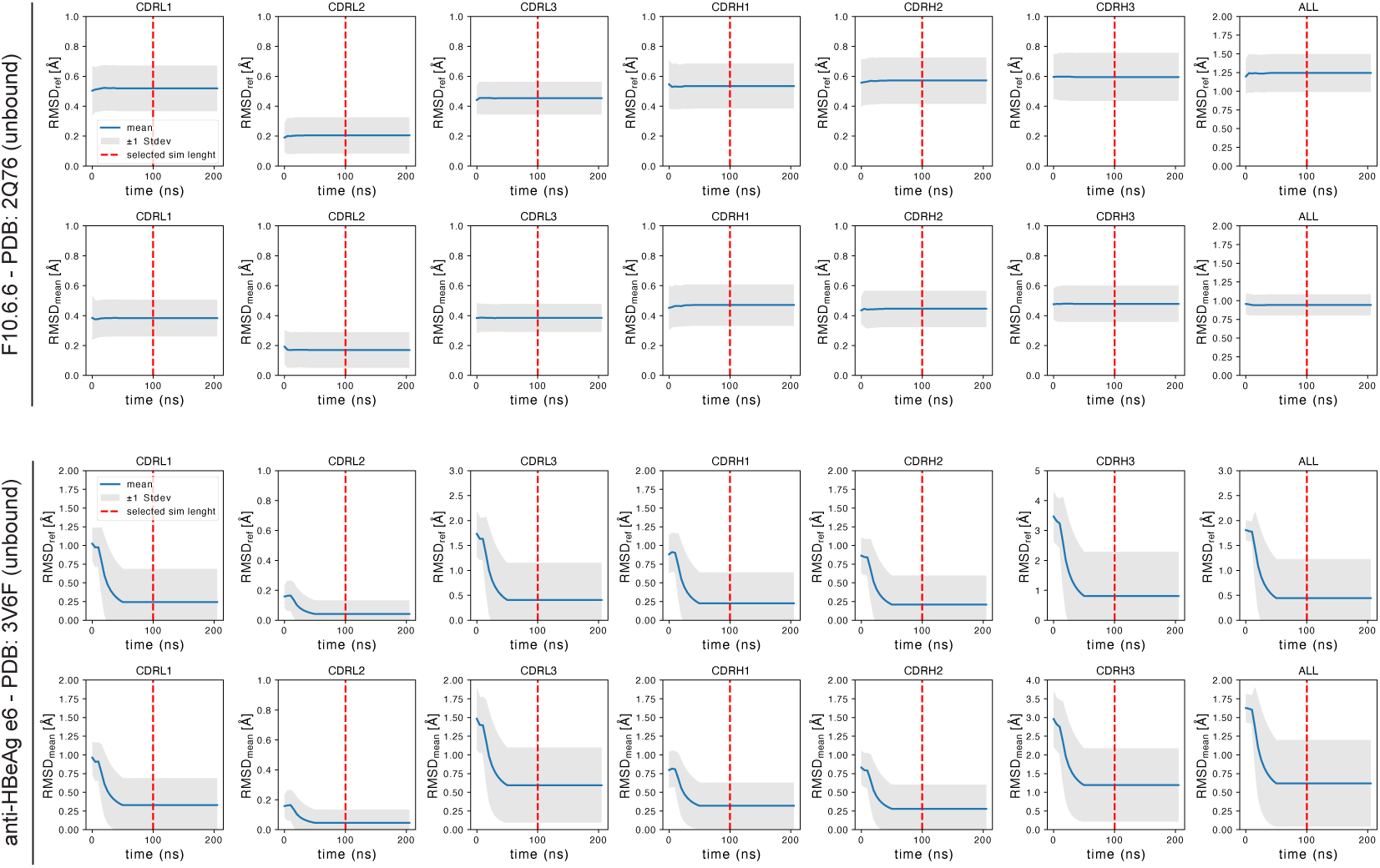
Convergence curves for coarse-grained simulations of benchmark antibodies. For each antibody system, the upper panels show the cumulative mean RMSD relative to the starting structure for each CDR, while the lower panels show the cumulative mean RMSD relative to the mean coordinates. Each simulation replica is represented by a different colour, and shaded areas indicate the standard deviation around the cumulative mean. The vertical red threshold line shows the final simulation length selected for all simulated systems.

